# A functional screen of translated pancreatic lncRNAs identifies a microprotein-independent role for *LINC00261* in endocrine cell differentiation

**DOI:** 10.1101/2020.04.28.062679

**Authors:** Bjoern Gaertner, Sebastiaan van Heesch, Valentin Schneider-Lunitz, Jana Felicitas Schulz, Franziska Witte, Susanne Blachut, Steven Nguyen, Regina Wong, Ileana Matta, Norbert Hubner, Maike Sander

## Abstract

Long noncoding RNAs (lncRNAs) are a heterogenous group of RNAs, which can encode small proteins. The extent to which developmentally regulated lncRNAs are translated and whether the produced microproteins are relevant for human development is unknown. Here, we show that many lncRNAs in direct vicinity of lineage-determining transcription factors (TFs) are dynamically regulated, predominantly cytosolic, and highly translated during pancreas development. We genetically ablated ten such lncRNAs, most of them translated, and found that nine are dispensable for endocrine cell differentiation. However, deletion of *LINC00261* diminishes generation of insulin^+^ endocrine cells, in a manner independent of the nearby TF *FOXA2*. Systematic deletion of each of *LINC00261*’s seven poorly conserved microproteins shows that the RNA, rather than the microproteins, is required for endocrine development. Our work highlights extensive translation of lncRNAs into recently evolved microproteins during human pancreas development and provides a blueprint for dissection of their coding and noncoding roles.

**Graphical Abstract:** 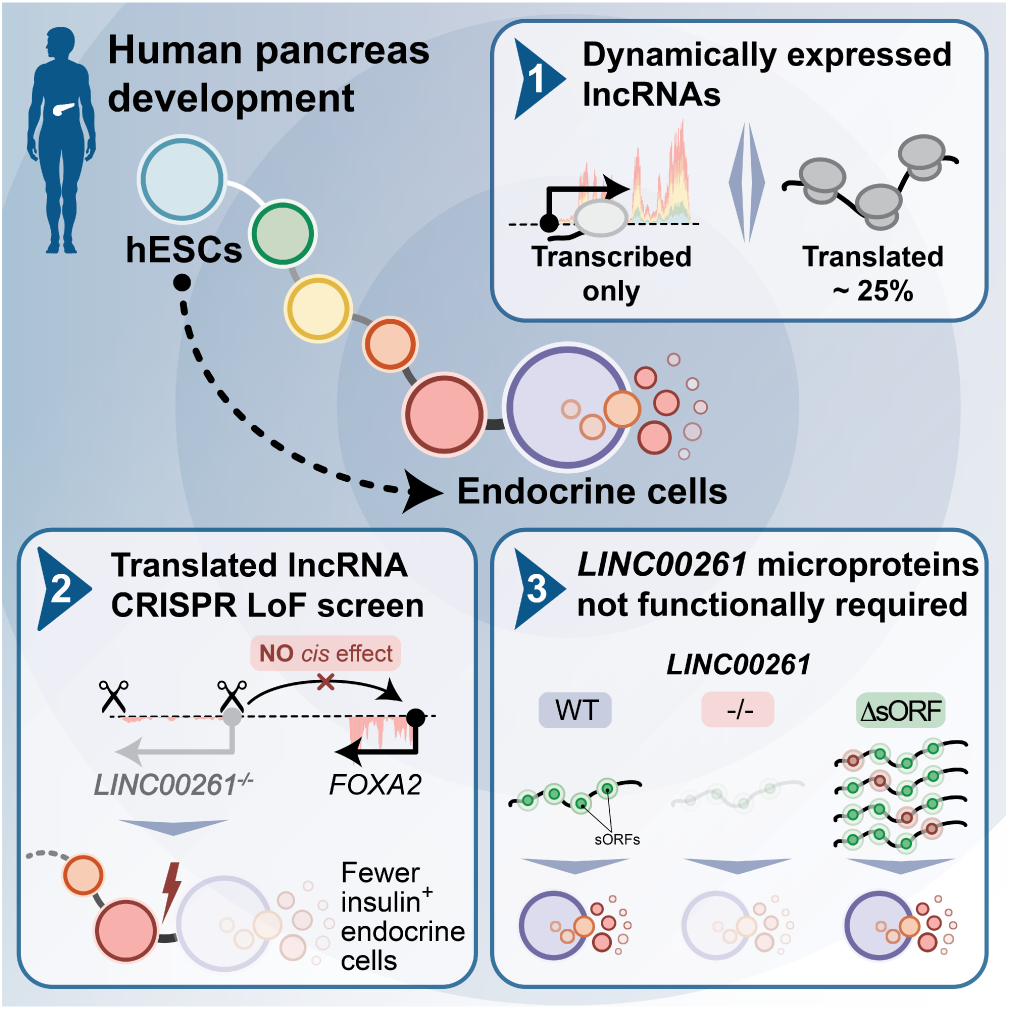

**Highlights:**

- Extensive lncRNA translation and microprotein production during human pancreas development
- A small-scale loss-of-function screen shows most translated lncRNAs are dispensable
- *LINC00261* is highly translated and regulates endocrine cell differentiation
- Deleting *LINC00261*’s evolutionary young microproteins reveals no essential roles

## INTRODUCTION

Defects in pancreatic endocrine cell development confer increased diabetes risk later in life (Bakhti et al., 2019). Therefore, a detailed understanding of the factors that orchestrate endocrine cell differentiation is highly relevant to human disease. Many of the molecular mechanisms that underlie the formation of pancreatic endocrine cells have been defined (Schiesser & Wells, 2014; Romer & Sussel, 2015), but a thorough functional assessment of the noncoding transcriptome, and in particular that of long noncoding RNAs (lncRNAs), is currently lacking.

Most lncRNAs with to date demonstrated roles in the regulation of fundamental developmental processes are active in the cell’s nucleus (Klattenhoff et al., 2013; Lin et al., 2014; Jiang et al., 2015; Kurian et al., 2015; Ramos et al., 2015; Daneshvar et al., 2016; Luo et al., 2016). However, a large proportion of lncRNAs is predominantly cytosolic (van Heesch et al., 2014; Cabili et al., 2015), and the functional relevance of these lncRNAs has remained unexplored in the context of human development. It is now widely accepted that many cytosolic lncRNAs possess short, “non-canonical” open reading frames (sORFs) that are actively translated (Bazzini et al., 2014; Ruiz-Orera et al., 2014; Makarewich & Olson, 2017). Although most of these non-canonical ORFs produce microproteins that are poorly conserved across species, recent studies have systematically assessed their biological activity, revealing roles across cellular organelles and, for a subset of microproteins, essential functions for cell survival (van Heesch et al., 2019; Chen et al., 2020; Prensner et al., 2020). This previously unrecognized coding capacity of supposedly noncoding RNAs has called into question the noncoding classification of lncRNAs, emphasizing the need for careful dissection of any gene’s coding and noncoding functions.

LncRNAs, whether translated or fully noncoding, are not randomly distributed in the genome but frequently located close to, and coregulated with, canonical protein-coding genes in *cis* (Luo et al., 2016; Neumann et al., 2018; van Heesch et al., 2019). For example, the lncRNAs *DIGIT* (also known as *GSC-DT*) and *Gata6as* (also known as *lncGata6* or *GATA6-AS1*) have been reported to enhance expression of the divergently expressed endoderm regulators *Goosecoid* (*GSC*) and *Gata6*, respectively (Daneshvar et al., 2016 ; Luo et al., 2016; Neumann et al., 2018). Similarly, *LINC00261* (also known as *DEANR1*) and its neighboring transcription factor (TF) *FOXA2* are both induced in endoderm formation, during which *LINC00261* has been demonstrated to positively regulate FOXA2 expression (Jiang et al., 2015). However, whether such *cis*-acting lncRNAs are translated and may exert cytosolic functions through *trans*-acting, microprotein-dependent mechanisms relevant for endoderm and pancreas development is not known.

In this study, we newly identify actively translated lncRNAs and analyze their role in human pancreas development. We accomplished this by classifying lncRNAs based on their dynamic regulation, subcellular localization, and active translation during the stepwise differentiation of human embryonic stem cells (hESCs) toward the pancreatic fate. We used this classification to prioritize select dynamically regulated and highly translated lncRNAs for deletion in hESCs, followed by extensive phenotypic characterization across multiple intermediate stages of pancreas development. This small-scale loss-of-function screen reveals that nine out of the ten selected lncRNAs are not essential for pancreatic development and, despite their vicinity to lineage-determining TFs, none of these lncRNAs regulate the expression of these TFs in *cis*.

The deletion of one lncRNA, *LINC00261*, does impair human endocrine cell development and leads to a significant reduction in the number of insulin-producing cells. Contrary to previous studies of *LINC00261* knockdown hESCs (Jiang et al., 2015), deletion of *LINC00261* has no effect on the expression of nearby TF *FOXA2* or other proximal genes, suggesting control of endocrine cell formation through a *trans*- rather than *cis*-regulatory mechanism. *LINC00261* is one of the most highly translated lncRNAs based on ribosome profiling (Ribo-seq) and produces multiple microproteins with distinct subcellular localizations. To systematically assess *LINC00261*’s coding and noncoding functions, we separately introduced frame shift mutations into each of seven identified *LINC00261* sORFs. However, rigorous phenotypic characterization revealed no apparent consequences of loss of each of the seven *LINC00261*-sORF-encoded microproteins on endocrine cell development. Our comprehensive assessment of functional lncRNA translation identifies a microprotein-independent *trans*-regulatory role for *LINC00261* in endocrine cell differentiation and provides a blueprint for the proper dissection of a gene’s coding and noncoding roles in a human disease-relevant system.

## RESULTS

### LncRNAs and nearby lineage-determining transcription factors exhibit dynamic coregulation during pancreas development

To identify lncRNAs involved in the regulation of pancreas development, we profiled RNA expression at five defined stages of hESC differentiation toward the pancreatic lineage: hESCs (ES), definitive endoderm (DE), primitive gut tube (GT), early pancreatic progenitor (PP1), and late pancreatic progenitor (PP2) (**Figure 1A**). While some lncRNAs were constitutively expressed (n = 592; 25.3%), the majority showed dynamic expression patterns, being either strongly enriched in (n = 874; 37.4%), or specific to (n = 871; 37.3%) a single developmental intermediate of pancreatic lineage progression (**Figure 1B** and **Table S1A**). The expression of many of these dynamically regulated lncRNAs correlated with that of proximal coding genes (**Figure S1A-D** and **Table S1B**,**C**), further exemplified by a subset of lncRNAs that was specifically coregulated with the key endodermal and pancreatic TFs *GATA6, FOXA2, PDX1*, and *SOX9* (**Figure 1C**,**D**). The tight expression coregulation of these lncRNA-TF pairs is likely explained by a shared chromatin environment (**Figure S1E-H**), which raises the possibility that like the TFs, the function of the lncRNAs is also required for endoderm and pancreas development.

**Figure 1.**
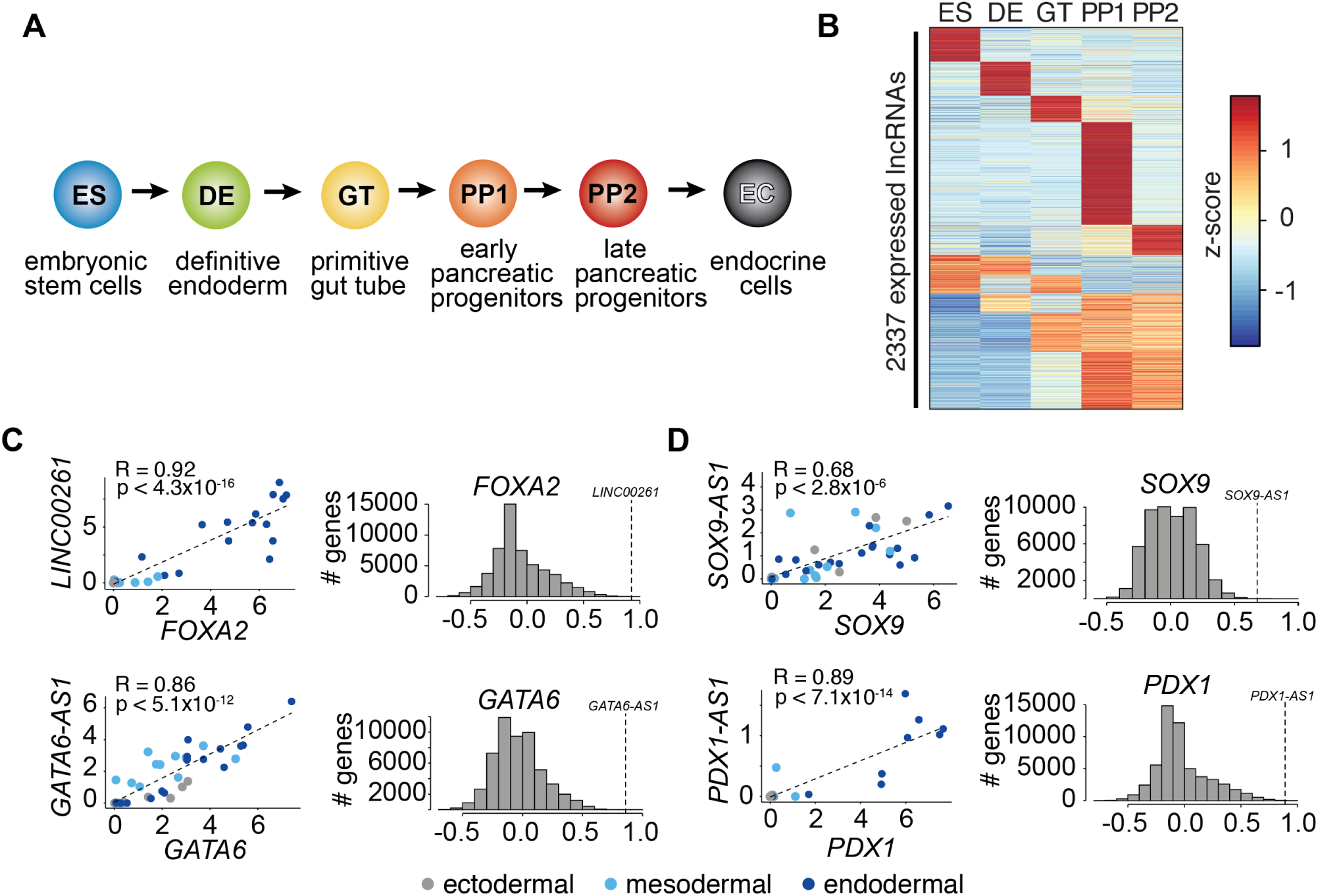
LncRNA expression and regulation during pancreatic differentiation. **(A)** Stages of directed differentiation from human embryonic stem cell (hESCs) to hormone-producing endocrine cells. The color scheme for each stage is used across all figures. **(B)** K-means clustering of all lncRNAs expressed (RPKM ≥ 1) during pancreatic differentiation based on their expression z-score (mean of n = 2 independent differentiations per stage; from CyT49 hESCs). Ten clusters were required (k = 10). **(C**,**D)** Left: Scatterplots comparing the expression of early (C) and late (D) expressed endodermal transcription factors (TFs) with the expression of their neighboring lncRNAs across 38 tissues. The dot color indicates the germ layer of origin of these tissues. Pearson correlation coefficients and p-values (t-test) are displayed. Right: Distribution of the Pearson correlation coefficients for each TF with all Ensembl 87 genes across the same 38 tissues. Dashed lines denote the correlation for the neighboring lncRNA, which for all lncRNAs shown is higher than expected by chance. See also **Figure S1** and **Table S1**.

### Many pancreatic progenitor-expressed lncRNAs are cytoplasmically enriched and translated

Although most functional roles described for lncRNAs to date have been predominantly nuclear (Marchese et al., 2017), multiple recent studies have shown that many lncRNAs are cytosolic and actively translated into sometimes biologically active microproteins (Makarewich & Olson, 2017). To further characterize the above-identified dynamically regulated lncRNAs, we analyzed their subcellular localization and translation potential using fractionation RNA-seq and Ribo-seq (**Figure 2A**). Of all lncRNAs expressed in pancreatic progenitor cells (PP2 cells), we classified 21% (n = 347) as localized to the nucleus, whereas a larger number (n = 563; 34%) primarily resided in the cytosol (**Figure S2A** and **Table S2A**). This subcellular distribution of pancreatic lncRNAs is in agreement with previous lncRNA localization studies by us and others (Clark et al., 2012; van Heesch et al., 2014; Cabili et al., 2015; Sun et al., 2015). LncRNAs enriched in the cytosol were expressed at higher levels than nucleus-localized lncRNAs, with expression levels similar to canonical protein-coding mRNAs (**Figure S2B**). Intriguingly, almost half of all cytosol-enriched lncRNAs (278 out of 563; 49.4%) displayed dynamic expression regulation during the differentiation of hESCs to pancreatic progenitors, suggesting that many lncRNAs with putative developmental functions do not act in the nucleus, but instead in the cytosol where they may be translated.

**Figure 2.**
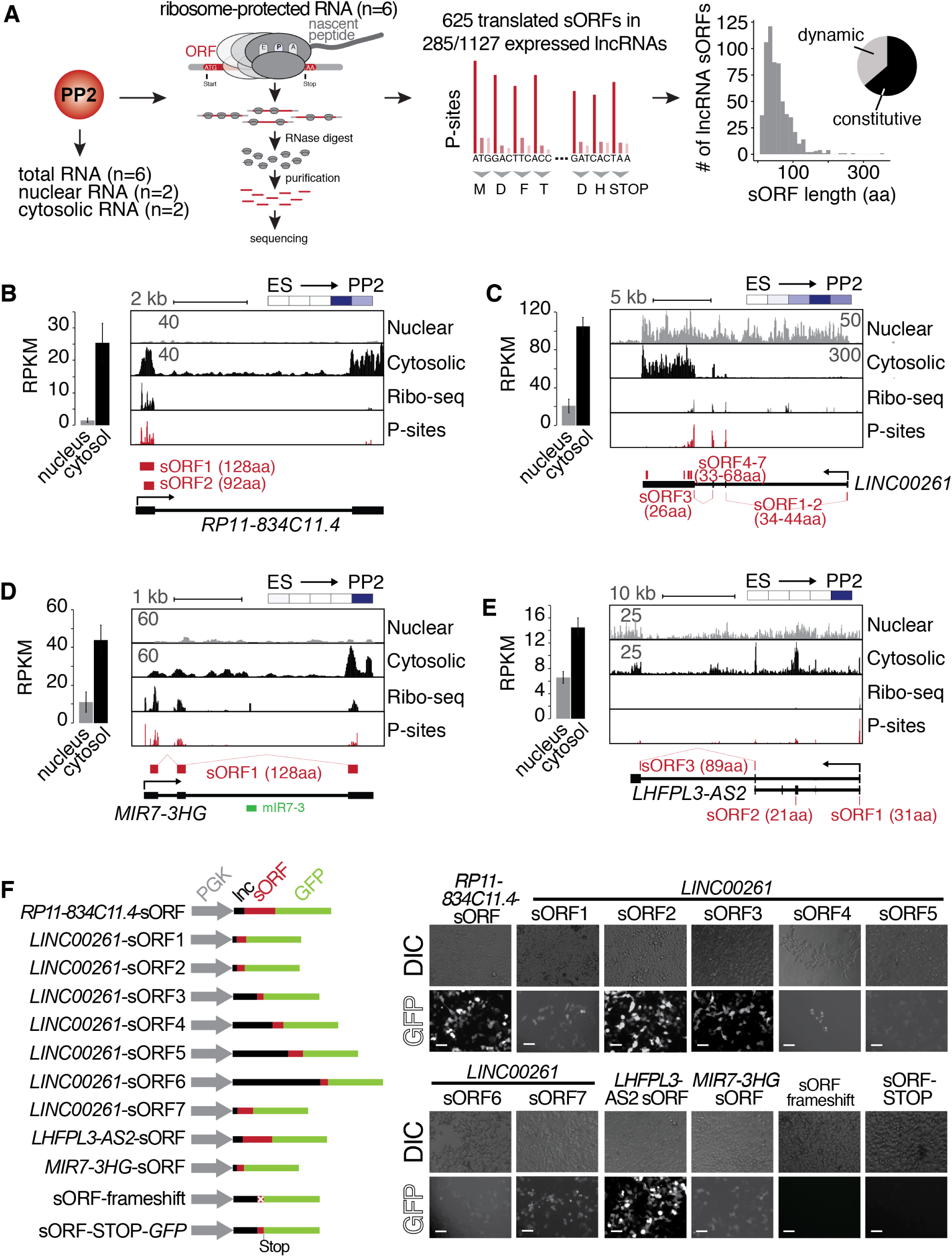
Cytosolic lncRNAs contain translated small open reading frames. **(A)** Overview of experimental strategy for subcellular fractionation and Ribo-seq-based identification of translated small open reading frames (sORFs) from lncRNAs expressed in PP2 cells. Replicates from six independent differentiations to the PP2 stage each for total (polyA) RNA-seq and Ribo-seq experiments, and two biological replicates for the subcellular fractionation were analyzed. The histogram on the far right depicts the size distribution of the sORF-encoded small peptides as number of amino acids (aa). The pie chart summarizes the percentages of constitutively and dynamically expressed sORF-encoding lncRNAs during pancreatic differentiation of CyT49 hESCs. **(B-E)** Left: Bar graphs showing nuclear and cytosolic expression (in RPKM) of lncRNAs *RP11-834C11*.*4* (B), *LINC00261* (C), *MIR7-3HG* (D), and *LHFPL3-AS2* (E). Data are shown as mean *±* S.D. (n = 2 biological replicates). Right: Subcellular fractionation RNA-seq, Ribo-seq, and P-site tracks (ribosomal P-sites inferred from ribosome footprints on ribosome-protected RNA) for loci of the depicted lncRNAs. Identified highest stringency sORFs (ORF in 6/6 replicates) are shown in red. For *LINC00261*, visually identified sORFs 1 and 2 are also shown. Heatmaps in the top right visualize the relative expression of the shown lncRNAs during pancreatic differentiation (means of two biological replicates per stage), on a minimum (white)/maximum (dark blue) scale. **(F)** *In vivo* translation reporter assays testing whether sORFs computationally defined in (A) give rise to translation products in HEK293T cells when fused in-frame to a GFP reporter. Left: Schematic of the constructs (gray: *PGK* promoter, black: lncRNA sequence 5’ to sORF to be tested, red: sORF, green: GFP ORF). Right: Representative DIC and GFP images of HEK293T cells transiently transfected with the indicated reporter constructs. Scale bars = 50 µm. See also **Figure S2** and **Table S2**.

To investigate the translation potential of these cytosolic lncRNAs, we used Ribo-seq, obtaining exceptionally deep and high quality translatome coverage across six replicate differentiations (**Figure S2C** and **Table S2B**). As nearly 90% of the sequenced ribosomal footprints exhibited clear 3-nucleotide codon movement characteristic of active translation (**Figure S2D-F**), these data have strong predictive value for the computational detection of non-canonical ORFs in lncRNAs (**Table S2C**). Requiring stringent reproducibility criteria (the exact ORF needed to be detected by RiboTaper (Calviello et al., 2016) in at least four out of the six replicates), we identified a total of 625 new sORFs in lncRNAs with a median length of 47 amino acids (aa) (**Table S2D**). The majority of detected sORFs (76%; n = 477/625) is currently not present in the sORFs.org database (Olexiouk et al., 2016) and thus completely novel. The translated sORFs are located within 285 cytosolically localized lncRNAs (25.3% of all expressed lncRNAs) (**Figure S2B**), which are expressed at higher levels than untranslated lncRNAs (**Figure S2G**) and exhibit translational efficiencies similar to mRNAs (**Figure S2H** and **Table S2E**).

Of note, only few of the newly identified sORFs are highly conserved across species, as judged by their low PhyloCSF scores (Lin et al., 2011) (**Table S2D**). However, the relevance of our ORF detection approach and the importance of lowly conserved ORFs for human biology have recently been demonstrated by several independent studies, focusing on either cardiac biology (van Heesch et al., 2019) or human cancer cell survival (Chen et al., 2020; Prensner et al., 2020).To our knowledge, our data constitute the first comprehensive set of non-canonical human ORFs generated from a non-transformed human cell model of development, providing a valuable resource for future functional studies.

### Translated pancreatic lncRNAs produce microproteins with distinct subcellular localizations

Having established that many stage-specific pancreatic lncRNAs are actively translated, we next sought to validate their translation potential through independent experimental approaches, and to demonstrate production of the predicted microproteins at the protein level. To this end, we first performed coupled *in vitro* transcription:translation assays on endogenous and complete transcript isoforms of four of the most highly translated lncRNAs (*LINC00261, RP11-834C11*.*4, LHFPL3-AS2*, and *MIR7-3HG*; **Figure S2I**; expression and ORF information in **Figure 2B-E**). Second, we generated a series of *in vivo* translation reporter constructs to assess the subcellular localization of microproteins translated from each of ten sORFs derived from the same four lncRNAs. Transient expression of individual constructs carrying in-frame GFP fusions in HEK293T cells produced GFP signal for all ten assayed microproteins, which was abolished upon introduction of a frameshift within the sORF or a stop codon following the sORF sequence (**Figure 2F** and **Figure S2J-L**). To rule out a possible localization bias induced by the GFP fusion, we also expressed a FLAG-tag fusion peptide (*RP11-834C11*.*4* sORF-1xFLAG), which revealed a cytoplasmic localization identical to the one observed for the GFP construct (**Figure S2J**). While most sORF-GFP fusion products were ubiquitously distributed throughout transfected cells, *LINC00261* sORF4-GFP specifically localized to mitochondria (**Figure S2K**), and *LINC00261* sORF7-GFP exhibited a perinuclear accumulation pattern reminiscent of aggresomes (**Figure S2L**). Taken together, our results validate the translation potential of sORFs encoded by pancreatic progenitor-expressed lncRNAs and show that these translation events result in robust production of microproteins with different subcellular localizations.

### Systematic knockout of translated lncRNAs during pancreas development

To identify potential functional roles of translated lncRNAs and the microproteins they produce in pancreas development, we selected ten candidates for CRISPR/Cas9-based genome editing in hESCs through excision of the lncRNA promoter or entire lncRNA locus (**Figure 3A**,**B**). These ten lncRNAs were prioritized based on (i) high expression and endodermal tissue-specificity, (ii) dynamic regulation during pancreas development, (iii) abundant translation of sORFs, and (iv) proximity to TFs with known roles in endoderm and pancreas development. For seven of the selected lncRNAs, translation was highly abundant and reproducibly detected across Ribo-seq replicates: *LINC00617* (also known as *TUNAR*; (Lin et al., 2014)), *GATA6-AS1* (also known as *GATA6-AS*; (Neumann et al., 2018)), *LINC00261, RP11-834C11*.*4, SOX9-AS1, MIR7-3HG*, and *LHFPL3-AS2*. Although for two additional lncRNAs the translation potential could not be determined, they were nonetheless included because of a previously reported requirement for definitive endoderm formation (*DIGIT*, also known as *GSC-DT*) (Daneshvar et al., 2016) and genomic localization adjacent to the definitive endoderm TF *LHX1* (*RP11-445F12*.*1*, also known as *LHX1-DT*). Lastly, *LINC00479* was chosen as a non-translated control with expression dynamics and a subcellular localization similar to *LINC00261*. Of note, for each of the ten selected lncRNAs, we generated at least two independent hESC knockout (KO) clones and used different combinations of single guide RNAs where possible (**Table S3A**).

**Figure 3.**
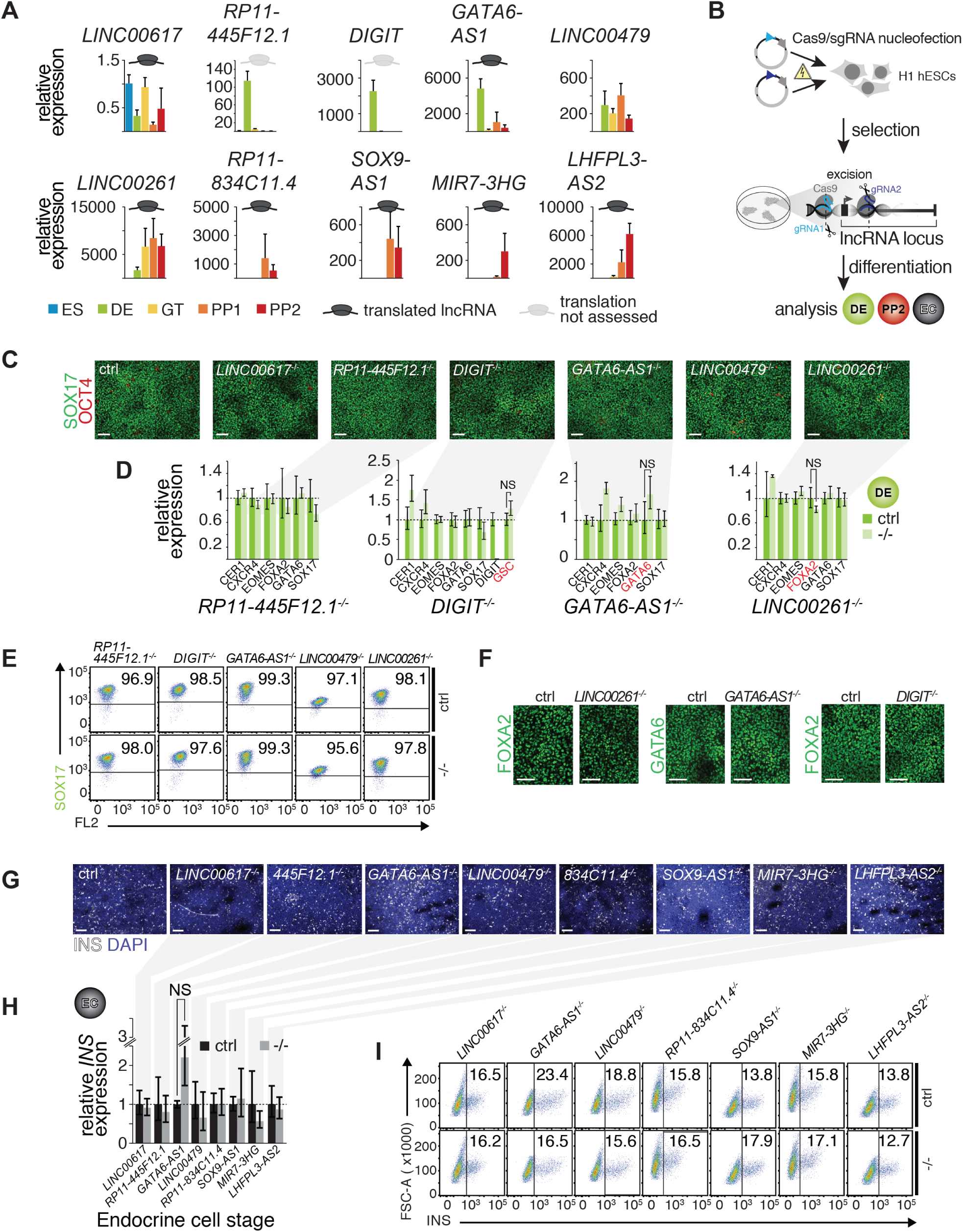
A small-scale CRISPR loss-of-function screen for dynamically expressed and translated lncRNAs during pancreatic differentiation. **(A)** qRT-PCR analysis of candidate lncRNAs during pancreatic differentiation of H1 hESCs relative to the ES stage. Data are shown as mean *±* S.E.M. (mean of n = 2-6 independent differentiations per stage; from H1 hESCs). **(B)** CRISPR-based lncRNA knockout (KO) strategy in H1 hESCs and subsequent phenotypic characterization. **(C)** Immunofluorescence staining for OCT4 and SOX17 in DE from control (ctrl) and KO cells for the indicated lncRNAs (representative images, n ≥ 3 independent differentiations; at least two KO clones were analyzed). **(D)** qRT-PCR analysis of DE lineage markers in DE from control and lncRNA KO (-/-) cells. TF genes in *cis* to the lncRNA locus are highlighted in red. Data are shown as mean *±* S.E.M. (n = 3-16 replicates from independent differentiations and different KO clones). NS, p-value > 0.05; t-test. **(E)** Flow cytometry analysis at DE stage for SOX17 in control and KO (-/-) cells for indicated lncRNAs. The line demarks isotype control. Percentage of cells expressing SOX17 is indicated (representative experiment, n ≥ 3 independent differentiations from at least two KO clones). **(F)** Immunofluorescence staining for FOXA2 or GATA6 in DE from control and *LINC00261, GATA6-AS1*, and *DIGIT* KO cells. **(G)** Immunofluorescence staining for insulin (INS) in endocrine cell stage (EC) from control and KO hESCs for the indicated lncRNAs (representative images, n ≥ 3 independent differentiations from at least two KO clones). **(H)** qRT-PCR analysis of INS in EC stage cultures from control and lncRNA KO (-/-) hESCs. Data are shown as mean *±* S.E.M. (n ≥ 4 replicates from independent differentiations of at least two KO clones). NS, p-value > 0.05; t-test. **(I)** Flow cytometry analysis at EC stage for INS in control and KO (-/-) cells for indicated lncRNAs. The line demarks isotype control. Percentage of cells expressing insulin is indicated (representative experiment, n ≥ 3 independent differentiations each from at least two KO clones). Scale bars = 100 µm. See also **Figure S3** and **Table S3**

We next differentiated each of the lncRNA KO hESC lines stepwise toward the pancreatic endocrine cell stage, conducting up to 16 replicate differentiations per clone. Because *LINC00617, RP11-445F12*.*1, DIGIT, GATA6-AS1, LINC00479*, and *LINC00261* were first expressed at, or before, the definitive endoderm stage (**Figure 3A**), we determined whether KO hESCs for these lncRNAs exhibited defects in definitive endoderm formation. Despite efficient lncRNA depletion (**Figure S3A**,**B**), neither quantification of definitive endoderm marker gene expression by qRT-PCR, nor immunofluorescence staining or flow cytometric analysis of the definitive endoderm marker SOX17 showed differences indicative of impaired endoderm formation in lncRNA KO lines (**Figure 3C-E**). Importantly, expression of TFs located in the direct vicinity of these lncRNAs, including *GSC* (*DIGIT*), *LHX1* (*RP11-445F12*.*1*), *GATA6* (*GATA6-AS1*), and *FOXA2* (*LINC00261*), was unaffected by the lncRNA KO (**Figure 3F, Figure S3C, Table S3B-D**), arguing against *cis*-regulation by these lncRNAs. These findings are in contrast to prior reports that have shown a requirement for *LINC00261* and *DIGIT* in definitive endoderm formation and the regulation of neighboring TFs *FOXA2* and *GSC*, respectively (Jiang et al., 2015; Daneshvar et al., 2016; Amaral et al., 2018; Swarr et al., 2019).

Next, we further differentiated control and KO lines for eight out of ten lncRNAs toward the endocrine cell stage, excluding *DIGIT* and *RP11-445F12*.*1* because they are not expressed after the definitive endoderm stage (**Figure 3A**). In KO hESC lines of seven out of these eight lncRNAs, we observed no effect on pancreatic progenitor cell formation or gene expression, with the exception of a few dysregulated genes in *LHFPL3-AS2* and *RP11-834C11*.*4* KO cells (**Figure S3C** and **Table S3E-K**). Furthermore, deletion of neither of the seven lncRNAs impaired endocrine cell formation, as determined by quantification of insulin^+^ cells and insulin mRNA levels (**Figure 3G-I**). Similar to the RNA expression results obtained at the definitive endoderm stage, deletion of none of the lncRNAs close to pancreatic TFs (e.g. *GATA6-AS1* and *SOX9-AS1*) altered the expression of these TFs, once more arguing against *cis*-regulation of these TFs by the neighboring lncRNA (**Figure S3C**).

Thus, nine out of ten endoderm- and pancreatic progenitor-enriched lncRNAs functionally investigated here appear to be nonessential for induction of the pancreatic fate and formation of insulin^+^ cells. Furthermore, these lncRNAs do not control the transcription of their proximal TFs.

### *LINC00261* knockout impairs endocrine cell development

The exception is the endoderm-specific lncRNA *LINC00261*, which is highly expressed and translated in pancreatic progenitors (**Figure S4A** and **Figure 2C**). While deletion of *LINC00261* caused no discernable phenotype in definitive endoderm (**Figure 3C-F** and**Figure S3C**), we observed a significant 30-50% reduction in the number of insulin^+^ cells at the endocrine cell stage (**Figure 4A**,**B**). This reduction in insulin^+^ cell numbers was consistent across four independently derived *LINC00261* KO hESC lines. In agreement with the reduced insulin^+^ cell numbers, insulin content and insulin mRNA levels were also reduced in *LINC00261* KO endocrine stage cultures (**Figure 4C**,**D**). Analysis of insulin fluorescence intensities by flow cytometry further showed no reduction in insulin levels per cell in one *LINC00261* KO clone and a mild reduction in the three other clones (**Figure 4E**), demonstrating that *LINC00261* predominately regulates endocrine cell differentiation rather than maintenance of insulin production in beta cells.

**Figure 4.**
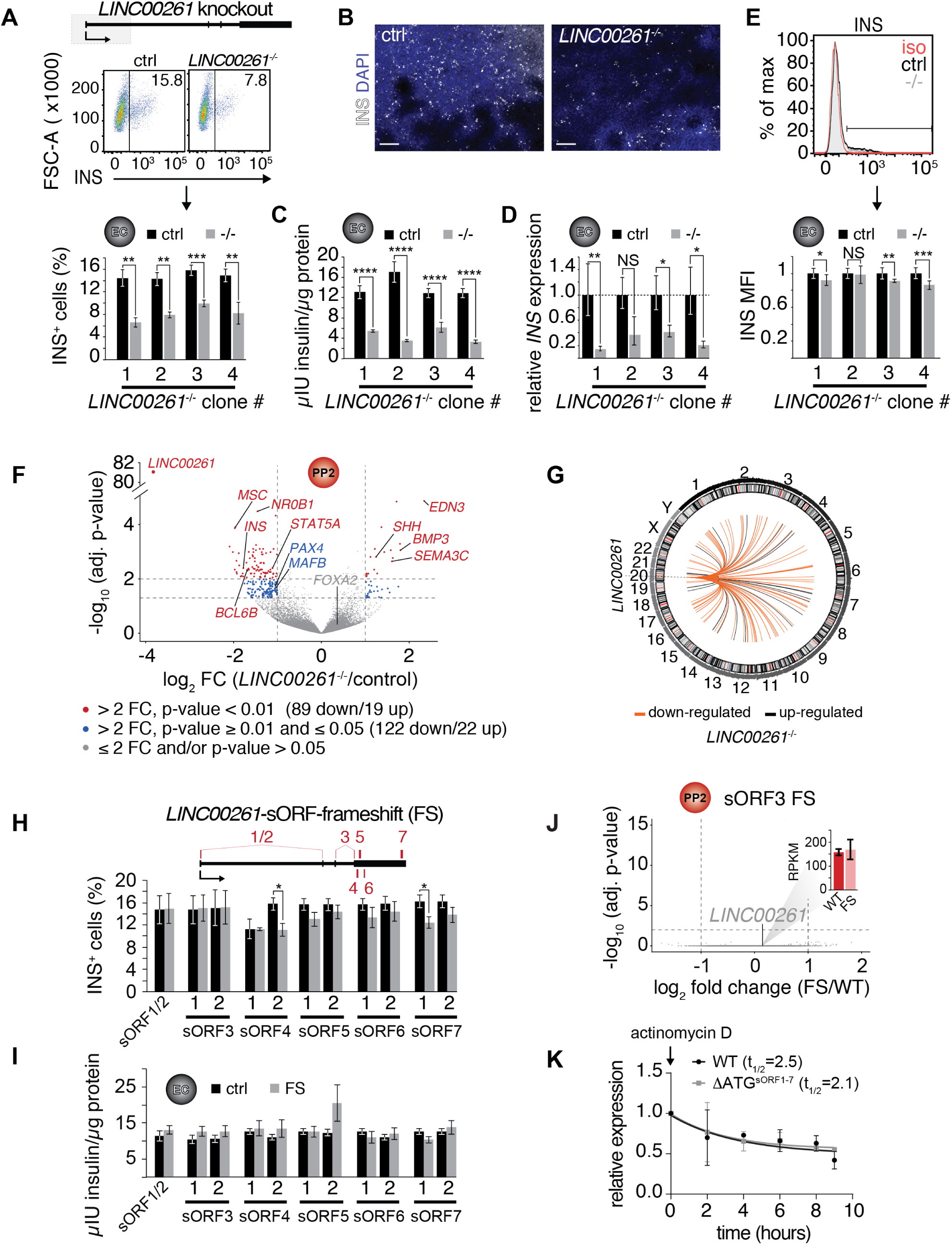
*LINC00261* deletion impedes pancreatic endocrine cell differentiation. **(A)** Flow cytometry analysis at endocrine cell stage (EC) for insulin (INS) in control (ctrl) and *LINC00261*^-/-^ H1 hESCs. Top panel: Schematic of the LINC00261 locus. The dashed box represents the genomic deletion. Middle panel: The line demarks isotype control. Percentage of cells expressing INS is indicated (representative experiment, n = 4 deletion clones generated with independent sgRNAs). Bottom panel: Bar graph showing percentages of INS-positive cells. Data are shown as mean *±* S.D. (n = 5 (clone 1), n = 6 (clone 2), n = 8 (clone 3), n = 5 (clone 4) independent differentiations). **(B)** Immunofluorescence staining for INS in EC stage cultures from control and *LINC000261*^-/-^ hESCs (representative images, number of differentiations see A). **(C)** ELISA for INS in EC stage cultures from control and *LINC00261*^-/-^ hESCs. Data are shown as mean *±* S.D. (n = 3 (clone 1), n = 2 (clone 2), n = 14 (clone 3), n = 13 (clone 4) independent differentiations). **(D)** qRT-PCR analysis of INS in EC stage cultures from control and *LINC00261*^-/-^ hESCs. Data are shown as mean *±* S.E.M. (n = 8 (clone 1), n = 5 (clone 2), n = 11 (clone 3), n = 3 (clone 4) independent differentiations). **(E)** Quantification of median fluorescence intensity after INS staining of control and *LINC00261*^-/-^ EC stage cultures. Data are shown as mean *±* S.D. (n = 5 (clone 1), n = 6 (clone 2), n = 8 (clone 3), n = 5 (clone 4) independent differentiations). iso, isotype control. **(F)** Volcano plot displaying gene expression changes in control versus *LINC00261*^-/-^ PP2 cells (n = 6 independent differentiations from all four deletion clones). Differentially expressed genes are shown in red (DESeq2; > 2-fold change (FC), adjusted p-value < 0.01) and blue (> 2-fold change, adjusted p-value ≥0.01 and ≤ 0.05). Thresholds are represented by vertical and horizontal dashed lines. *FOXA2* in *cis* to *LINC00261* is shown in gray (gray dots represent genes with ≤ 2-fold change and/or adjusted p-value > 0.05). **(G)** Circos plot visualizing the chromosomal locations of the 108 genes differentially expressed (DESeq2; > 2-fold change (FC), adjusted p-value < 0.01) in *LINC00261*^-/-^ compared to control PP2 cells, relative to *LINC00261* on chromosome 20. No chromosome was over- or underrepresented (Fisher test, p-value > 0.05 for all chromosomes). **(H)** Top panel: Schematic of the *LINC00261* locus, with the location of its sORFs (1 to 7) marked by vertical red bars. Bottom panel: Flow cytometric quantification of INS-positive cells in control and *LINC00261*-sORF-frameshift (FS) at the EC stage. Data are shown as mean *±* S.D. (n = 4-7 independent differentiations per clone). **(I)** ELISA for INS in EC stage cultures from control and *LINC00261*-sORF-FS hESCs. Data are shown as mean *±* S.D. (n = 3-7 independent differentiations per clone). **(J)** Volcano plot displaying gene expression changes in control versus *LINC00261*-sORF3-FS PP2 cells. No gene was differentially expressed (DESeq2; > 2-fold change, adjusted p-value < 0.01; indicated by dashed horizontal and vertical lines; n = 2 independent differentiations). *LINC00261* is shown in gray, the bar graph insert displays *LINC00261* RPKM values in control and *LINC00261*-sORF3-FS PP2 cells. **(K)** *LINC00261* half-life measurements in HEK293T cells transduced with lentivirus expressing either wild type (WT) *LINC00261* or ΔATG^sORF1-7^ *LINC00261* (mutant in which the ATG start codons of sORFs 1-7 were changed to non-start codons). HEK293T were treated with the transcription inhibitor actinomycin D and RNA isolated at 0, 2, 4, 6, 8, and 9 hours post actinomycin D addition. *LINC00261* expression was analyzed by qRT-PCR relative to the TBP gene. Data are shown as mean *±* S.E.M. (n = 3 biological replicates for each assay time point). *, p-value < 0.05; **, p-value < 0.01; ***, p-value < 0.001; ****, p-value < 0.0001; NS, p-value > 0.05; t-test. Scale bars = 100 µm. See also **Figure S4** and **Table S4**.

To determine the molecular effects of *LINC00261* deletion, we performed RNA-seq in pancreatic progenitors derived from *LINC00261* KO and control hESCs. Among the down-regulated genes were the TFs *MAFB* and *PAX4* (**Figure 4F, Figure S4B, Table S4A**), which are important regulators of beta cell differentiation (Sosa-Pineda et al., 1997; Artner et al., 2007). Similar to the absence of *cis*-regulatory functions observed in the other lncRNA KOs, we found no evidence for *cis*-regulation of *FOXA2* by *LINC00261* (**Figure 4F** and **Figure S4C**). Of note, the genes differentially expressed in *LINC00261* KO cells mapped to all chromosomes and showed no enrichment for chromosome 20 where *LINC00261* resides (**Figure 4G**). Combined, our results suggest a *trans*- rather than *cis*-regulatory function for *LINC00261*, consistent with its predominantly cytosolic localization, active translation, and diffuse distribution within the nucleus (**Figure 2C** and **Figure S4D**). This potential *trans* functionality prompted us to further investigate whether *LINC00261*’s coding or noncoding features are essential for endocrine cell differentiation.

### The *LINC00261* transcript, and not the encoded microproteins, is required for endocrine cell differentiation

We established that *LINC00261* harbors multiple distinct and highly translated sORFs, which produce poorly conserved microproteins with diverse subcellular localizations (**Figure 2C**,**F, Figure S2I**,**K**,**L, Table S2D**). This raises the possibility that *LINC00261*-sORF-encoded microproteins, and not the RNA itself, are functionally important for endocrine cell differentiation. To systematically dissect whether the microproteins are required for endocrine cell formation independent of *LINC00261* RNA, we individually mutated all seven sORFs in independent hESC lines, leaving the lncRNA sequence grossly intact. Each of these hESC lines either carries a homozygous frameshift mutation near the microprotein’s N-terminus (for sORFs 1-6) or a full sORF deletion (sORF7; **Table S4B**). After verifying that CRISPR editing of the *LINC00261* locus did not impact *LINC00261* transcript levels (**Figure S4E**), we quantified (i) insulin mRNA levels, (ii) insulin^+^ cells, and (iii) total insulin content in endocrine cell stage cultures. We observed no difference between sORF loss-of-function and control hESC lines for any of these endpoints (**Figure 4H**,**I** and **Figure S4E**). Consistently, transcriptome analysis of pancreatic progenitors with frameshifts in sORF3 (the most highly translated *LINC00261*-sORF; **Figure 2C** and **Table S2D**) revealed no differentially expressed genes between *LINC00261*-sORF3 frameshift and control cells (**Figure 4J** and **Table S4C**), contrasting observations in *LINC00261* RNA KO pancreatic progenitors (**Figure 4F** and **Table S4A**).

It has been suggested that ribosome association can degrade lncRNAs, e.g. through nonsense-mediated decay (Tani et al., 2013; Carlevaro-Fita et al., 2016). Therefore, to determine whether the multiple sORFs within *LINC00261* regulate *LINC00261* stability, we simultaneously mutated start codons of all seven sORFs (ΔATG^sORF1-7^ *LINC00261*) and expressed either wild type or ΔATG^sORF1-7^ *LINC00261* ectopically in HEK293T cells where *LINC00261* is normally not expressed. *LINC00261* half-life measurements upon transcriptional inhibition with actinomycin D revealed no difference in *LINC00261* levels between wild type and ΔATG^sORF1-7^ *LINC00261* (**Figure 4K**), suggesting that the association of *LINC00261* with ribosomes does not affect its stability.

In sum, through the systematic, one-by-one removal of microproteins produced from a highly translated lncRNA with functional importance for pancreatic endocrine cell formation, we found no evidence to implicate the individual microproteins in endocrine cell development. Although *LINC00261*’s microproteins may share functional redundancy or have developmental roles that do not affect the production of insulin^+^ cells, our findings strongly suggest that by themselves, each of the *LINC00261*-sORF-encoded microproteins is not functionally required for endocrine cell formation.

## DISCUSSION

### The highly translated lncRNA *LINC00261* is a novel regulator of endocrine cell differentiation

In this study, we globally characterized molecular features of lncRNAs expressed during progression of hESCs toward the pancreatic lineage, including their subcellular localization and potential to be translated and to produce microproteins. We performed a phenotypic CRISPR loss-of-function screen, focusing on ten developmentally regulated, highly expressed, and highly translated lncRNAs proximal to TFs known to regulate pancreas development.

The first important observation from this screen is that we find no evidence to implicate the lncRNAs *LINC00261, DIGIT, GATA6-AS1, SOX9-AS1*, and *RP11-445F12*.*1* in the *cis*-regulation of their neighboring TFs *FOXA2, GSC, GATA6, SOX9*, and *LHX*, respectively, despite tight transcriptional coregulation of the lncRNA-TF pairs.

Second, we identify the translated lncRNA *LINC00261* as a novel regulator of pancreatic endocrine cell differentiation, as evidenced by a severe reduction in insulin^+^ cell numbers upon *LINC00261* deletion. We show that *LINC00261* transcripts are highly abundant in pancreatic progenitors and, albeit present in the nucleus, are predominantly localized to the cytoplasm. Here, they frequently associate with ribosomes to produce multiple distinct microproteins. Through the introduction of individual frameshift mutations in each of the microprotein-encoding sORFs of *LINC00261*, we could uncouple the requirement of *LINC00261* in endocrine cell development from microprotein production. Furthermore, mutating all translated *LINC00261* sORFs simultaneously and thereby significantly reducing *LINC00261*’s ability to bind ribosomes, did not affect *LINC00261* transcript levels.

This indicates that, in contrast to some reports suggesting that translated sORFs can regulate RNA stability by promoting nonsense-mediated RNA decay (Tani et al., 2013; Carlevaro-Fita et al., 2016), the high translation levels and multiple sORFs of *LINC00261* are not part of a *LINC00261* decay pathway.

Although lncRNAs are now appreciated as a novel and abundant source of sORF-encoded biologically active microproteins (Makarewich & Olson, 2017), we found no essential roles for *LINC00261*-sORF-encoded microproteins in endocrine cell development. Possibly, this is explained by the fact that most microproteins - including the majority of microproteins newly identified in this study, and all microproteins produced by *LINC00261* - are poorly conserved across species. Since the functional role of the vast majority of such recently evolved microproteins has not been systematically investigated, there is an ongoing debate about their significance for vital cellular processes (Ruiz-Orera et al., 2018; Levy, 2019). Recent reports, however, suggest that lowly conserved sORFs can indeed have important functions in terminally differentiated cells and in cancer (van Heesch et al., 2019; Chen et al., 2020; Prensner et al., 2020). This raises the possibility that *LINC00261*-sORF-encoded microproteins could possess functions that become relevant under specific environmental, developmental, or disease conditions not examined in this study.

### *LINC00261* - a potential *trans* regulator of endocrine cell differentiation?

Several lines of evidence suggest that *LINC00261* regulates endocrine cell differentiation in *trans*: (i) *LINC00261* transcripts show a diffuse distribution in multiple subcellular compartments, (ii) genes differentially expressed in *LINC00261* KO cells are randomly distributed throughout the genome, (iii) expression of the nearby TF *FOXA2* is not affected by *LINC00261* deletion. Such a *trans* regulatory mechanism for *LINC00261* is supported by a recent study from the GTEx Consortium, where *LINC00261* is highlighted as one of a few lncRNAs that forms a potential *trans* regulatory hotspot through genetic interactions that influence the expression of multiple distant genes (Aguet et al., 2019). Consistent with its preferential cytosolic localization, and further supporting the notion of a *trans* regulatory mechanism, *LINC00261* has been suggested to regulate gene expression through non-nuclear mechanisms, e.g. by preventing nuclear translocation of *β*-catenin (Wang et al., 2017) or by acting as a miRNA sponge (Shi et al., 2019; Wang et al., 2019; Yan et al., 2019). Although our observations and current literature strongly hint to a function in *trans* independent of the produced microproteins, the exact mechanism by which *LINC00261* regulates gene expression in pancreatic progenitors remains to be determined.

We here present a rigorous, in-depth characterization of dynamically regulated and translated lncRNAs in a disease-relevant cell context of human developmental progression. Our combination of ultra-high-coverage RNA and Ribo-seq, protein-level validation of microprotein production and localization, and the systematic deletion of all individual microproteins encoded by a single translated lncRNA, not only provides a detailed resource of translated ‘non-canonical’ sORFs and their microproteins in pancreatic development, but also serves as a blueprint for the systematic functional interrogation of translated lncRNAs during human organ development.

## Acknowledgements

We thank Andrea Carrano for comments on the manuscript and Francesca Mulas for advice with computational analyses. We acknowledge the UCSD IGM Genomics Center for next generation sequencing (P30 DK063491) and the UCSD Human Embryonic Stem Cell Core Facility for assistance with flow cytometry analysis and cell sorting. This work was supported by the National Institutes of Health (R01 DK068471 and R01 DK078803 to M.S.), an Alexander von Humboldt Foundation Research Award to M.S., and a postdoctoral fellowship from the Larry L. Hillblom Foundation (2015-D-021-FEL to B.G.). S.v.H. was supported by an EMBO long-term fellowship (ALTF 186-2015, LTFCOFUND2013, GA-2013-609409). N.H. is the recipient of an ERC advanced grant under the European Union Horizon 2020 Research and Innovation Program (grant agreement AdG788970).

## Author Contributions

B.G. and M.S. conceived the project, and B.G., M.S., S.v.H., and N.H. designed the experiments. B.G., S.v.H., J.S., S.B., S.N., R.W., and I.M. performed experiments. B.G., V.S.-L., and F.W. conducted bioinformatics analysis. B.G., M.S., S.v.H., and N.H. interpreted data. B.G., S.v.H., and M.S. wrote the manuscript.

## Materials and Methods

### HEK293T cell culture

HEK293T cells (female) were cultured in a humidified incubator at 37 °C with 5% CO_2_ using Dulbecco’s Modified Eagle Medium (Cat# 45000-312; 4.5 g/L glucose, [+] L-glutamine, [-] sodium pyruvate) supplemented with 10% fetal bovine serum (FBS).

### hESC culture and maintenance

H1 hESCs (male) were grown in feeder-independent conditions on

Matrigel®-coated dishes (Corning) with mTeSR1 media (STEMCELL Technologies). Propagation was carried out by passing the cells every 3 to 4 days using Accutase™ (eBioscience) for enzymatic cell dissociation. hESC research was approved by the University of California, San Diego, Institutional Review Board and Embryonic Stem Cell Research Oversight Committee.

### Pancreatic differentiation

H1 hESCs were differentiated in a monolayer format as previously described (Rezania et al., 2012), with minor modifications. Undifferentiated hESCs were seeded into 24-wells at 0.4 *×* 10^6^ cells/well in 500 *µ*l mTeSR1 medium. The next day the cells were washed in RPMI media (Thermo Fisher Scientific) and then differentiated with daily media changes. In addition to GlutaMAX™, RPMI medium was supplemented with 0.12% (w/v) NaHCO_3_ and 0.2% (Day 0) or 0.5% (Day 1-3) (v/v) FBS (Corning). DMEM/F12 medium (Corning; 45000-350) was supplemented with 2% (v/v) FBS and 0.2% (w/v) NaHCO_3_, and DMEM High Glucose medium (HyClone) was supplemented with 0.5X B-27™ supplement (Thermo Fisher Scientific).

Human Activin A, mouse Wnt3a, human KGF, and human Noggin were purchased from R&D Systems. Other media components included TGF*β* R1 kinase inhibitor IV (EMD Bioscience), KAAD-Cyclopamine (Toronto Research Chemicals), the retinoid analog TTNPB (Sigma Aldrich), the protein kinase C activator TPB (EMD Chemicals), the BMP type 1 receptor inhibitor LDN-193189 (Stemgent), and an inhibitor of the TGF-*β* type 1 activin like kinase receptor ALK5, ALK5 inhibitor II (Enzo Life Sciences).

Stage 1 (DE; collection on day 3):

Day 0: RPMI/FBS, 100 ng/mL Activin A, 25 ng/mL mouse Wnt3a

Day 1 – 2: RPMI/FBS, 100 ng/mL Activin A

Stage 2 (GT; collection on day 6):

Day 3: DMEM/F12/FBS, 2.5 *µ*M TGF*β* R1 kinase inhibitor IV, 50 ng/mL KGF

Day 4 – 5: DMEM/F12/FBS, 50 ng/mL KGF

Stage 3 (PP1; collection on day 10):

Day 6 – 9: DMEM/B27, 3nM TTNPB, 0.25 mM KAAD-Cyclopamine, 50 ng/mL Noggin

Stage 4 (PP2; collection on day 13):

Day 10 – 12: DMEM/B27, 100 nM ALK5 inhibitor II, 100 nM LDN-193189, 500 nM TPB, 50 ng/mL Noggin

Stage 5 (endocrine cell stage; collection on day 16):

Day 13 – 15: DMEM/B27, 100 nM ALK5 inhibitor II, 100 nM LDN-193189, 500 nM TPB, 50 ng/mL Noggin

For ribosome profiling experiments, a scalable suspension culture protocol was employed for differentiation of H1 cells to the PP2 stage (Rezania et al., 2014). Undifferentiated hESCs were aggregated by preparing a single cell suspension in mTeSR1 media (STEMCELL Technologies; supplemented with 10 *µ*M Y-27632) at 1 *×* 10^6^ cells/mL and overnight culture in six-well ultra-low attachment plates (Costar) with 5.5 ml per well on an orbital rotator (Innova2000, New Brunswick Scientific) at 100 rpm. The following day, undifferentiated aggregates were washed in MCDB 131 media (Thermo Fisher Scientific) and then differentiated using a multistep protocol with daily media changes and continued orbital rotation at either 100 rpm or at 115 rpm from days 8 to 14. In addition to 1% GlutaMAX™ (Gibco) and 10 mM (days 0-10) or 20 mM (days 11-14) glucose, MCDB 131 media was supplemented with 0.5% (days 0-5) or 2% (days 6-14) fatty acid-free BSA (Proliant), 1.5 g/L (days 0-5 and days 11-14) or 2.5 g/L (days 6-10) NaHCO_3_ (Sigma-Aldrich), and 0.25 mM ascorbic acid (days 3-10). Human Activin A, mouse Wnt3a, and human KGF were purchased from R&D Systems. Other media components included Insulin-Transferrin-Selenium-Ethanolamine (ITS-X; Thermo Fisher Scientific; days 6-10), retinoic acid (RA) (Sigma-Aldrich), the sonic hedgehog pathway inhibitor SANT-1 (Sigma-Aldrich), the protein kinase C activator TPB (EMD Chemicals), the BMP type 1 receptor inhibitor LDN-193189 (Stemgent), and the TGF*β* type 1 activin like kinase receptor ALK5 inhibitor, ALK5 inhibitor II (Enzo Life Sciences).

Stage 1 (DE; collection on day 3):

Day 0: MCDB 131, 100 ng/mL Activin, 25 ng/mL mouse Wnt3a

Day 1 – 2: MCDB 131, 100 ng/mL Activin A

Stage 2 (GT; collection on day 6):

Day 3 – Day 5: MCDB 131, 50 ng/mL KGF

Stage 3 (PP1; collection on day 8)

Day 6 – Day 7: MCDB 131, 50ng/mL KGF, 0.25 *µ*M SANT-1, 1 µM RA 100 nM LDN-193189, 200 nM TPB

Stage 4 (PP2; collection on day 11):

Day 8 – Day 10: MCDB 131, 2ng/mL KGF, 0.25 *µ*M SANT-1, 0.1 µM RA, 200 nM LDN-193189, 100 nM TPB

### CRISPR/Cas9-mediated lncRNA knockout

To generate clonal lncRNA knockout hESC lines, combinations of pSpCas9(BB)-2A-Puro plasmid pairs

(Addgene plasmid # 62988, gift from Feng Zhang) expressing Cas9 and single sgRNAs targeting upstream and downstream regions of the lncRNA promoter/locus were co-transfected into 1.5 *×* 10^6^ H1 hESCs using the Human Stem Cell Nucleofector Kit 2 (Lonza) and the Amaxa Nucleofector II (Lonza). 24 hours after plating into Matrigel®-coated six-well plates, nucleofected cells were selected with puromycin (1 *µ*g/mL mTeSR1 media) for 2-3 consecutive days. Individual colonies that emerged within 7 days after transfection were subsequently transferred manually into 96-well plates for expansion. Genomic DNA for PCR genotyping with GoTaq® Green Mastermix (Promega) and Sanger sequencing was then extracted using QuickExtract™ DNA Extraction Solution (Lucigen). The *PDX1* knockout line was generated in an analogous way.

To generate sORF frameshift mutations, sgRNA sequences targeting the N-terminal region of the predicted small peptides were inserted into pSpCas9(BB)-2A-GFP (Addgene plasmid #48138, gift from Feng Zhang) via its BpiI cloning sites. 3 *µ*g of the resulting plasmids were then transfected into 500,000 H1 cells plated into Matrigel®-coated six-wells the day prior, using XtremeGene 9 Transfection Reagent (Sigma-Aldrich) according to the manufacturer’s instructions. 24 hours post-transfection, 10,000 GFP+ cells were sorted on an Influx™ Cell Sorter (BD Biosciences) into Matrigel®-coated six-wells containing 1 mL mTeSR1 media supplemented with 10 *µ*M ROCK inhibitor and 1X penicillin/streptomycin. Seven days after sorting, emerging colonies were hand-picked and transferred into 96-well plates for genotyping. Frame-shifts inside the targeted sORFs were confirmed by PCR-amplification of the sORF sequence with GoTaq® Green Mastermix and subsequent subcloning the PCR products into pCR2.1 (Thermo Fisher Scientific). For each hESC clone, at least six pCR2.1 clones were Sanger sequenced. Oligonucleotide sequences for sgRNA cloning are provided in **Table S5A**.

### PCR genotyping of CRISPR clones

Four days after transfer of single cell-derived clones into 96-wells, cell culture supernatants containing dead cells were collected from each well prior to the daily media change. Cell debris was then pelleted and used for gDNA extraction with 10-20 *µ*l QuickExtract™ DNA Extraction Solution (Lucigen) according to the manufacturer’s instructions. 1 *µ*l DNA was then PCR-amplified with GoTaq® Green Mastermix (Promega) and locus-specific primers that anneal either within or outside of the excised genomic DNA. PCR products generated with “inside” primers were visualized on a 2% agarose gel, PCR bands generated with primers flanking the deletion were gel-purified and submitted for Sanger sequencing (see **Table S5B** for genotyping and sequencing primers).

For genotyping of sORF frameshift clones, PCR amplicons designed to encompass the Cas9 cut site were amplified and Sanger sequenced (**Table S5B**). If out-of-frame indels were apparent in the sequencing chromatogram, the sequenced PCR product was ligated into pCR2.1-TOPO via TOPO-TA cloning. A minimum of six clones were Sanger sequenced in order to determine the genotype at both alleles with high confidence.

### Generation of sORF translation reporter plasmids

The four lncRNAs tested were PCR-amplified with KOD Xtreme™ DNA Hotstart Polymerase (Millipore) from their 5’ end up until the last codon of the sORF to be tested, omitting its stop codon (primer sequences are listed in **Table S5D**). cDNA was used as PCR template for *LINC00261* and *LHFPL3-AS2*; *RP11-834C11*.*4*, and *MIR7-3HG* were amplified from a gBlock synthetic gene fragment (Integrated DNA Technologies; see **Table S5F**). The GFP coding sequence (without start codon; amplified from pRRLSIN.cPPT.PGK-GFP.WPRE) was then fused in-frame to the sORF via overlap extension PCR. The resulting fusion product was cloned into pRRLSIN.cPPT.PGK-GFP.WPRE via BshTI and SalI restriction sites included in the PCR primers. Due to the 3’-location of sORF7 within *LINC00261*, not the entire *LINC00261* cDNA was amplified but only 65 bp preceding sORF7.

To create the *RP11-834C11*.*4*-sORF-1XFLAG reporter construct in an analogous way, a gBlock synthetic gene fragment encompassing the FLAG-tagged sORF served as PCR template (**Table S5F**). The resulting PCR product was cloned into pRRLSIN.cPPT.PGK-GFP.WPRE via BshTI and SalI restriction sites.

### Generation of *LINC00261* wild type and ΔATG^sORFS1-7^ expression plasmids

The *LINC00261* wild type cDNA was PCR-amplified from pENTR/D-TOPO-LINC00261 (gift from Leo Kurian) with KOD Xtreme™ DNA Hotstart Polymerase (Millipore). The resulting PCR product was inserted into pRRLSIN.cPPT.PGK-GFP.WPRE via its appended BshTI/SalI cloning sites. Full-length *LINC00261* ΔATG^sORFS1-7^ was assembled through overlap extension PCR from the following three fragments and subsequently cloned into pRRLSIN.cPPT.PGK-GFP. WPRE via appended BshTI/SalI cloning sites: (i) a 1,248 bp PCR product amplified from a synthetic gene construct (Genewiz; see **Table S5F** for sequence) in which the ATG start codons of sORFs 1-6 had been mutated (ATG AAG / ATT / AGG / AAG / ATA / AGG), and (ii-iii) 3,111 bp/610 bp PCR fragments (amplified from the *LINC00261* cDNA) in which the sORF7 start codon was mutated (ATG AAG). The obtained plasmids were sequence-verified by Sanger sequencing.

### Immunofluorescence staining

H1 hESC-derived cells grown as monolayer on Matrigel®-coated coverslips were washed twice with PBS and then fixed with 4% paraformaldehyde in PBS for 30 min at room temperature. Following three washes in PBS, samples on coverslips were permeabilized and blocked with Permeabilization/Blocking Buffer (0.15% (v/v) Triton X-100 and 1% normal donkey serum in PBS) for 1 hour at room temperature. Primary and secondary antibodies were diluted in Permeabilization/Blocking Buffer. Sections were incubated overnight at 4°C with primary antibodies, and then secondary antibodies for 30 min at room temperature. The following primary antibodies were used: rabbit anti-OCT4 (Cell Signaling Technology, 1:500), goat anti-SOX17 (Santa Cruz Biotechnology, 1:250), goat anti-FOXA2 (Santa Cruz Biotechnology, 1:250), goat anti-GATA6 (Santa Cruz Biotechnology, 1:50), guinea pig anti-insulin (Dako). Secondary antibodies (1:1000) were Cy3-, Cy5-, Alexafluor488-conjugated antibodies raised in donkey against guinea pig, rabbit, mouse, or goat (Jackson Immuno Research Laboratories). Images were acquired on a Zeiss Axio-Observer-Z1 microscope with a Zeiss AxioCam digital camera, and figures prepared with Adobe Photoshop/Illustrator CS5.

### Flow cytometry analysis

For intracellular flow cytometry, single cells were washed three times in FACS buffer (0.1% (w/v) BSA in DPBS) and then fixed and permeabilized with Cytofix/Cytoperm Fixation/Permeabilization Solution (BD Biosciences) for 20 min at 4 °C, followed by two washes in BD Perm/Wash™ Buffer. Cells were next incubated with either PE-conjugated anti-SOX17 antibody (BD Biosciences), or PE-conjugated anti-INS antibody (Cell Signaling Technology) in 50 µl BD Perm/Wash™ Buffer for 60 min at 4 °C. Following three washes in BD Perm/Wash™ Buffer, cells were analyzed on a FACSCanto II (BD Biosciences) cytometer.

### Insulin content measurements

To measure total insulin content of endocrine cell stage control and lncRNA KO cells, adherent cultures were enzymatically detached from a 24-well at day 16 of differentiation. Upon quenching with FACS buffer (0.1% (w/v) BSA in DPBS), the cells were pelleted and extracted over night at 4 °C in 100 µl acid-ethanol (2% HCl in 80% ethanol). Insulin was measured by Insulin ELISA (Alpco) and normalized to total protein, as quantified with a BCA protein assay (Thermo Fisher Scientific).

### Quantitative reverse transcription PCR (qRT-PCR)

Total RNA was isolated from hESC-derived cells and HEK293T cells using either TRIzol® (Thermo Fisher Scientific) or the RNAeasy Mini Kit (Qiagen), respectively. Upon removal of genomic DNA (TURBO DNA-free™ Kit or RNase-free DNase Set) cDNA was synthesized using the iScript™ cDNA Synthesis Kit (Bio-Rad). PCR reactions were run in triplicate with 6.25-12.5 ng cDNA per reaction using the CFX96 Real-Time PCR Detection System (BioRad). TATA-binding protein (TBP) was used as endogenous control to calculate relative gene expression using the ΔΔCt method. Primer sequences are provided in **Table S5C**.

### Transient transfection of HEK293T cells with polyethylenimine (PEI)

Two hours prior to transfection, fresh pre-warmed DMEM medium (Corning) was added to each well.

Transfection mix was prepared by combining PEI and plasmid DNA (4:1 ratio; 4 µg PEI per 1 *µ*g DNA) in Opti-MEM™ Reduced Serum Medium (Thermo Fisher Scientific) followed by brief vortexing. After five minutes, the transfection complex was added dropwise to the cells.

### Lentivirus preparation and ectopic *LINC00261* expression

Lentiviral particles were prepared by co-transfecting HEK293T cells (using PEI) with the pCMVR8.74/pMD2.G helper plasmids and with pRRLSIN.cPPT.PGK-GFP.WPRE transfer plasmid, in which the GFP ORF had been replaced with the 4.9 kb *LINC00261* cDNA. Virus-containing supernatant was collected for two consecutive days and concentrated by ultracentrifugation for 2 hours at 19,400 rpm using an Optima L-80 XP Ultracentrifuge (Beckman Coulter).

To express *LINC00261* (wild type) and *LINC00261* (ΔATG^sORF1-7^) in HEK293T cells, the cells were plated in 6-well plates and transduced with lentivirus the following day. Two days post infection, the cells were passaged for RNA half-life measurements

### *LINC00261* RNA half-life measurement

HEK293T cells transduced with either *LINC00261* (wild type) or *LINC00261* (ΔATG^sORF1-7^) lentivirus were seeded in six 24-wells. 48 hours after plating, cells from one well were collected for RNA isolation as the “0 hour” time point. To the remaining five wells, 100 *µ*l growth media supplemented with 10 *µ*g/ml actinomycin D were added to inhibit transcription. At 2, 4, 6, 8, and 9 hours following actinomycin D addition, samples were collected for RNA isolation. Total RNA was then reverse transcribed and analyzed by qPCR, where the abundance of each time point was calculated relative to the abundance at the 0 hour time point (ΔCt). The half-life was then determined by non-linear regression (One phase decay; GraphPad Prism).

### Single molecule RNA fluorescence *in situ* hybridization (smRNA FISH)

H1-derived PP2 stage cells (control and LINC00261 KO) were cultured on Matrigel®-coated 12 mm diameter coverslips in a 24-well plate. Following 10 min fixation in 1 mL Fixation Buffer (3.7 % (v/v) formaldehyde in 1X PBS) at room temperature, the cells were washed twice in PBS and subsequently permeabilized in 70 % (v/v) ethanol for one hour at 4 °C. Following a five minute wash in Stellaris RNA FISH Wash Buffer A (LGC Biosearch Technologies; 1:5 diluted concentrate, with 10% (v/v) formamide added), the coverslips were incubated in a humidified chamber at 37 °C for 14 hours with probes diluted in Stellaris RNA FISH Hybridisation Buffer (LGC Biosearch Technologies; with 10% (v/v) formamide added) to 125 nM. After a 30 min wash at 37 °C in Wash Buffer A, the cells were counter-stained with Hoechst 33342 (Thermo Fisher Scientific) for 15 min and washed in RNA FISH Wash Buffer B (LGC Biosearch Technologies) for 5 min at room temperature. The coverslips were mounted in Vectashield Mounting Medium (Vector Laboratories) and imaged on a UltraView Vox Spinning Disk confocal microscope (PerkinElmer) using a 100X oil objective.

### *In vitro* transcription/translation of lncRNAs

Synthetic gene constructs containing complete transcript isoforms (including the predicted 5’ and 3’ UTR) of four translated lncRNAs (*RP11-834C11*.*4, LINC00261, MIR7-3HG*, and *LHFPL3-AS2*) were produced by Genewiz (constructs available upon request). Microproteins were translated *in vitro* from 0.5 *µ*g linearized plasmid DNA using the TnT® Coupled Wheat Germ Extract system (Promega) in the presence of 10 mCi/mL [35S]-methionine (Hartmann Analytic) according to manufacturer’s instructions. 5 *µ*L lysate was denatured for 2 min at 85 °C in 9.6 *µ*L Novex Tricine SDS Sample Buffer (2X) (Thermo Fisher Scientific) and 1.4 µL DTT (500 mM). Proteins were separated on 16% Tricine gels (Thermo Fisher Scientific) for 1 h at 50 V followed by 3.5 h at 100 V and blotted on PVDF-membranes (Immobilon-PSQ Membrane, Merck Millipore). Incorporation of [35S]-methionine into newly synthesized proteins enabled the detection of translation products by phosphor imaging (exposure time of 1 day).

### *In vivo* translation assays

Reporter plasmids were transfected into HEK293T cells using PEI, and 36 hours post transfection live cells were imaged on an EVOS Cell Imaging System (Thermo Fisher Scientific) equipped with a 20X objective. Additional constructs were generated that served as negative controls (no GFP fluorescence):

1) a *LINC00261*-sORF3-GFP construct with a single ‘T’ insertion inside sORF3, causing a frame-shift,

2) a *LINC00261*-sORF2-GFP construct with a stop codon preceding the GFP coding sequence, and

3) a *LINC00261*-sORF1-GFP construct with a frame-shift mutation within the GFP coding sequence.

### Stranded mRNA-seq library preparation for lncRNA KOs

Total RNA from PP2 cells differentiated with the Rezania et al. (2012) protocol was isolated and DNase-treated using either TRIzol® (Thermo Fisher Scientific), or the RNAeasy Mini kit (Qiagen) according to the manufacturer’s instructions. RNA integrity (RIN >8) was verified on the Agilent 2200 TapeStation (Agilent Technologies), and 400 ng RNA was used for multiplex library preparation with the KAPA mRNA HyperPrep Kit (Roche). All libraries were evaluated on TapeStation High Sensitivity DNA ScreenTapes (Agilent Technologies) and with the Qubit dsDNA High Sensitivity (Life Technologies) assays for size distribution and concentration prior to pooling the multiplexed libraries for single-end 1×51nt or 1×75 sequencing on the HiSeq 2500 or HiSeq 4000 System (Illumina). Libraries were sequenced to a depth of >20M uniquely aligned reads.

### Cell fractionation and ribo-minus RNA-seq

H1 hESCs were differentiated to the PP2 stage with the Rezania et al. (2012) protocol, then nuclear and cytosolic RNA was isolated with the Paris™ Kit (Thermo Fisher Scientific). Unfractionated total RNA was set aside as a control. All samples were DNaseI-treated prior to further processing (TURBO DNA-free™ Kit; Thermo Fisher Scientific). rRNA-depleted total RNA-seq libraries were prepared with TruSeq® Stranded Total RNA Library Prep Gold (Illumina), and sequencing was performed on a HiSeq4000 instrument.

### Alignment of lncRNA KO mRNA-seq samples and processing for gene expression analysis

Using the Spliced Transcripts Alignment to a Reference (STAR) aligner (STAR 2.5.3b; (Dobin et al., 2013)), sequence reads were mapped to the human genome (hg38/GRCh38) with the Ensembl 87 annotations in 2-pass mapping mode, allowing for up to 6 mismatches. Cufflinks (part of the Cufflinks version 2.2.1 suite (Trapnell et al., 2010; Roberts et al., 2011)), was then used to quantify the abundance of each transcript cataloged in the Ensembl 87 annotations in reads per kilobase per million mapped reads (RPKM). For plotting expression values, a pseudocount of 1 was added to all RPKM values prior to log_2_-transformation.

Genes with RPKM ≥ 1 across two replicates were deemed expressed. Differential gene expression was tested using the DESeq2 v1.10.1 Bioconductor package (Love et al., 2014) with default parameters. Input count files for DESeq2 were created with htseq-count from the HTSeq Python library (Anders et al., 2015). Genes with a >2-fold change and an adjusted p-value of <0.01 were considered differentially expressed. The chromosomal localization of genes differentially expressed upon *LINC00261* KO was visualized with the RCircos package in R (https://cran.r-project.org/web/packages/RCircos/index.html).

### LncRNA classifications

The following transcript biotypes were grouped into the “lncRNA” classification: 3’ overlapping ncrna, antisense, bidirectional promoter lncRNA, lincRNA, macro lncRNA, non coding, processed transcript, sense intronic, sense overlapping, TEC.

LncRNAs with ≥ 1 RPKM during all differentiation stages of CyT49 hESCs (ES, DE, FG, GT, PP1, PP2) were categorized as constitutively expressed (“constitutive”), whereas lncRNAs with <1 RPKM throughout differentiation were considered “never expressed”. LncRNAs expressed in at least one of the stages (but not in all five stages) were referred to as dynamically expressed (“dynamic”). Furtheremore, for each lncRNA, its maximum RPKM value was determined across 38 tissues/cell types (see “Gene-gene correlations and GO enrichment” section below). Log_2_-transformed maximum expression values (RPKM + pseudocount of 1) were graphed as boxplots for different gene sets using the ggplot2 R package (https://cran.r-project.org/web/packages/ggplot2/index.html). To determine the subcellular localization of lncRNAs, first all lncRNAs expressed in the nuclear and/or cytosolic RNA fraction (RPKM ≥ 1 in two biological replicates) of H1-derived PP2 stage cells were selected. Among these expressed lncRNAs, those with ≥ 1 RPKM_cytosol_ and <1 RPKM_nucleus_ were classified as “cytosol enriched”. Conversely, lncRNAs with <1 RPKM_cytosol_ and ≥ 1 RPKM_nucleus_ were termed “nucleus enriched”. LncRNAs expressed in both fractions (≥ 1 RPKM_cytosol_ and ≥ 1 RPKM_nucleus_) were tagged with “both”.

### Assignment of lncRNAs to their nearest coding gene using GREAT

GREAT (Genomic Regions Enrichment of Annotations Tool 3.0.0; (McLean et al., 2010)) was run with the “Single nearest gene” within 1000 kb option to assign the nearest coding genes to the following sets of lncRNAs: i) DE-transcribed lncRNAs, ii) PP2-transcribed lncRNAs that are not transcribed at the DE stage (non-transcribed control set for i)), iii) PP2-transcribed lncRNAs, and iv) lncRNAs transcribed at the DE stage but not transcribed in PP2 cells (non-transcribed control set for iii)). The log_2_-transformed RPKM values of the lncRNA-associated coding genes were then graphed as boxplots using ggplot2. The corresponding absolute coding-to-lncRNA inter-gene distances were visualized as cumulative frequency plots.

### Gene-gene correlations and GO enrichment

Pearson correlations were calculated among all genes across a catalog of 38 tissues/cell types derived from all three germ layers (16 Illumina BodyMap 2.0 tissues, other publicly available data sets (see “Data Sources” below), and EndoC-*β*H1 RNA-seq data generated in our lab). Scatter plots of the log_2_-transformed RPKM values for lncRNAs/neighboring TFs and histograms of the Pearson correlation coefficients were plotted in R using ggplot2.

Spearman correlations were calculated to test for expression coregulation among all genes expressed (RPKM ≥ 1) in a minimum of ten out of 38 tissues. The resulting correlation matrix was then used to calculate the Euclidean distance followed by hierarchical clustering. The resulting heatmap was subdivided into ten clusters. Cluster visualization was done using heatmap.3 (https://raw.githubusercontent.com/obigriffith/biostar-tutorials/master/Heatmaps/heatmap.3.R) from gplots v3.0.1 (http://cran.r-project.org/web/packages/gplots/index.html). GO enrichment (Ashburner et al., 2000; The Gene Ontology, 2019) and KEGG pathway (Kanehisa et al., 2017) analyses to assign functional annotation to all ten clusters were performed with gProfiler v0.6.4 (Reimand et al., 2016) using g:Profiler archive revision 1741 (Ensembl 90, Ensembl Genomes 38).

### Alignment and processing of ChIP-seq samples

All sequence reads were filtered to include only those passing the standard Illumina quality filter, and then aligned to the Homo sapiens reference genome (hg38/GRCh38) using Bowtie version 1.1.1 (Langmead et al., 2009). The following parameters were used to select only uniquely aligning reads with a maximum of two mismatches:

-k 1 –m 1 –l 50 –n 2 –best –strata

SAMtools (Li et al., 2009) was then used to filter reads with a MAPQ score less than 30 and to remove duplicate reads. Finally, replicate ChIP-seq and input BAM files were merged and sorted. The HOMER makeUCSCfile function (Heinz et al., 2010) was used to create a bedGraph formatted file for viewing in the UCSC Genome Browser.

### Ribosome profiling and matching RNA-seq

Ribosome profiling was performed on PP2 cells obtained from six independent differentiations of H1 hESCs with the Rezania et al. (2014) protocol, yielding an average of 89% PDX1-positive cells. Ribosome footprinting and sequencing library preparation was done with the TruSeq® Ribo Profile (Mammalian) Library Prep Kit (Illumina) according to the TruSeq® Ribo Profile (Mammalian) Reference Guide (version August 2016). In short, 50 mg of PP2 aggregates were washed twice with cold PBS and lysed for 10 minutes on ice in 1mL lysis buffer (1 x TruSeq Ribo Profile mammalian polysome buffer, 1 % Triton X-100, 0.1% NP-40, 1 mM dithiothreitol, 10 U ml-1 DNase I, cycloheximide (0.1 mg/ml) and nuclease-free H_2_O). Per sample, 400 *µ*L of lysate was further processed according to manufacturer’s instructions. Final library size distributions were checked on the Bioanalyzer 2100 using a High Sensitivity DNA assay (Agilent Technologies), multiplexed and sequenced on an Illumina HiSeq 4000 producing single end 1×51 nt reads. Ribo-seq libraries were sequenced to an average depth of 85M reads.

Total RNA was isolated using TRIzol® Reagent (Thermo Fisher Scientific) from the exact same cell cultures processed for ribosome profiling (10% of the total number of cells). Total RNA was DNase treated and purified using the RNA Clean & Concentrator™-25 kit (Zymo Research). RIN scores (RIN = 10 for all 6 samples) were measured on a BioAnalyzer 2100 using the RNA 6000 Nano assay (Agilent Technologies). Poly(A)-purified mRNA-seq library preparation was performed according to the TruSeq Stranded mRNA Reference Guide (Illumina), using 500 ng of total RNA as input. Libraries were multiplexed and sequenced on an Illumina HiSeq 4000 producing paired-end 2×101nt reads.

### Alignment of Ribo-seq and matched mRNA-seq samples

Prior to mapping, ribosome-profiling reads were clipped for residual adapter sequences and filtered for mitochondrial, ribosomal RNA and tRNA sequences (**Table S2**). Next, all mRNA and ribosome profiling data were mapped to the Ensembl 87 transcriptome annotation of the human genome hg38 assembly using STAR 2.5.2b (Dobin et al., 2013) in 2-pass mapping mode. To avoid mRNA-seq mapping biases due to read length, 2×101 nt mRNA-seq reads were next trimmed to 29-mers, those mRNA reads were processed and mapped with the exact same settings as the ribosome profiling data. For the mapping of 2×101 nt RNA-seq reads 6 mismatches per read were allowed (default is 10), whereas 2 mismatches were permitted for the Ribo-seq and trimmed mRNA-seq reads. To account for variable ribosome footprint lengths, the search start point of the read was defined using the option *–seedSearchStartLmaxOverLread*, which was set to 0.5 (half the read, independent of ribosome footprint length). Furthermore, -*-outFilterMultimapNmax* was set to 20 and *–outSAMmultNmax* to 1, which prevents the reporting of multimapping reads.

### Detecting actively translated reading frames

Canonical ORF detection using ribosome profiling data was performed with RiboTaper v1.3 (Calviello et al., 2016) with standard settings. For each sample, we selected only the ribosome footprint lengths for which at least 70% of the reads matched the primary ORF in a meta-gene analysis. Following the standard configuration of RiboTaper, we required ORFs to have a minimum length of 8aa, evidence from uniquely mapping reads and at least 21 P-sites. The final list of translation events was stringently filtered requiring the translated gene to have an average RNA RPKM ≥ 1 and to be detected as translated in all 6 profiled samples. Furthermore, we required the exact ORF to be detected independently in at least 4 out of 6 samples.

### Translational efficiency estimates

Translational efficiency (TE) estimations were calculated as the ratio of Ribo-seq over mRNA-seq DESeq2 normalized counts, yielding independent gene-specific TEs for each of the 6 individual replicate differentiations. For this, mRNA-seq and Ribo-seq based expression quantification was calculated for (annotated and newly detected) coding sequences (CDSs / ORFs) only, using RNA reads trimmed to footprint sizes as described above.

### Data sources

The following datasets used in this study were downloaded from the GEO and ArrayExpress repositories:

RNA-seq: Illumina BodyMap 2.0 expression data from 16 human tissues (GSE30611); polyA mRNA RNA-seq from BE2C (GSE93448), GM12878 (GSE33480), 293T (GSE34995), HeLa (GSE33480), HepG2 (GSE90322), HUVEC (GSE33480), Jurkat (GSE93435), K562 (GSE33480), MiaPaCa-2 (GSE43770), Panc1 (GSE93450), PFSK-1 (GSE93451), U-87 MG (GSE90176); CyT49 hESC/DE/GT/PP1/PP2/CD142+ progenitors/CD200+ polyhormonal cells/in vivo matured endocrine cells/pancreatic islets (E-MTAB-1086). ChIP-seq: H3K4me3/H3K27me3 in CyT49 hESC/DE/GT/PP1/PP2 (E-MTAB-1086).

## Statistical analyses

Statistical analyses were performed using Microsoft Excel, GraphPad Prism (7.05), and R (v.3.5.0). Statistical parameters such as the value of n, mean, standard deviation (S.D.), standard error of the mean (S.E.M.), significance level (*p < 0.05, **p < 0.01, ***p < 0.001 and ****p < 0.0001), and the statistical tests used are reported in the figures and figure legends. The “n” refers to the number of independent pancreatic differentiation experiments analyzed (biological replicates), or the number of genes/transcripts and sORFs detected. Statistically significant gene expression changes were determined with DESeq2.

## Data availability

All mRNA-seq and Ribo-seq datasets generated for this study have been deposited at GEO under the accession number GSE144682.

## Supplemental Information

**Title: A functional screen of translated pancreatic lncRNAs identifies a microprotein-independent role for *LINC00261* in endocrine cell differentiation**

**Inventory:**

### I. Supplemental Data

**Figure S1**, Related to Figure 1

**Figure S2**, Related to Figure 2

**Figure S3**, Related to Figure 3

**Figure S4**, Related to Figure 4

**Table S1 - Related to Figure 1. Identification, regulation, and characterization of lncRNAs during pancreatic differentiation. (A)** Gene expression during pancreatic differentiation (RPKM). **(B)** lncRNA-proximal TFs, by cluster in correlation heatmap (Figure S1H). **(C)** GO enrichment and KEGG pathway analysis for each cluster in the correlation heatmap (Figure S1D).

(supplied as Excel file: Table S1.xlsx)

**Table S2 - Related to Figure 2. RNA-seq after subcellular fractionation and Ribo-seq in PP2 cells. (A)** Subcellular fractionation of PP2 stage cells (RPKM). **(B)** Ribo-seq/mRNA-seq contaminant filtering statistics, read size distribution, and Pearson correlation coefficients of all sequenced Ribo-seq and polyA RNA-seq libraries. **(C)** All ORFs detected by RiboTaper, including lncRNA sORFs. **(D)** lncRNA sORFs detected by RiboTaper and conservation statistics (PhyloCSF scores). **(E)** Translation efficiency calculations.

(supplied as Excel file: Table S2.xlsx)

**Table S3 - Related to Figure 3. Differentially expressed genes after lncRNA deletion. (A)** Coordinates of CRISPR deletions. **(B)** Differentially expressed genes in *RP11-445F12*.*1* knockout at definitive endoderm stage. **(C)** Differentially expressed genes in *GATA6-AS1* knockout at definitive endoderm stage. **(D)** Differentially expressed genes in *LINC00261* knockout at definitive endoderm stage. **(E)** Differentially 75 expressed genes in *LINC00617* knockout at PP2 stage. **(F)** Differentially expressed genes in *GATA6-AS1* knockout at PP2 stage. **(G)** Differentially expressed genes in *LINC00479* knockout at PP2 stage. **(H)** Differentially expressed genes in *RP11-834C11*.*4* knockout at PP2 stage. **(I)** Differentially expressed genes in *SOX9-AS1* knockout at PP2 stage. **(J)** Differentially expressed genes in *MIR7-3HG* knockout at PP2 stage. **(K)** Differentially expressed genes in *LHFPL3-AS2* knockout at PP2 stage.

(supplied as Excel file: Table S3.xlsx)

**Table S4 - Related to Figure 4. Characterization of *LINC00261* knockout and *LINC00261*-sORF3-frameshift PP2 cells. (A)** Differentially expressed genes in *LINC00261* knockout PP2 stage cells. **(B)** Sequences of *LINC00261* wild type and frameshift mutants. **(C)** Differentially expressed genes in *LINC00261*-sORF3-frameshift PP2 stage cells. (supplied as Excel file: Table S4.xlsx)

**Table S5 List of oligonucleotides and synthetic gene fragments used in this study. (A)** sgRNA oligonucleotides used for cloning into PX458/Px459. **(B)** Genotyping and sequencing primers for KO validation. **(C)** qRT-PCR primers. **(D)** Cloning primers (translation reporter constructs and lentiviral LINC00261 overexpression plasmids). **(E)** Synthetic gene fragments. **(F)** Custom *LINC00261* Stellaris® RNA FISH probe set. (supplied as Excel file: Table S5.xlsx)

**Figure S1.**
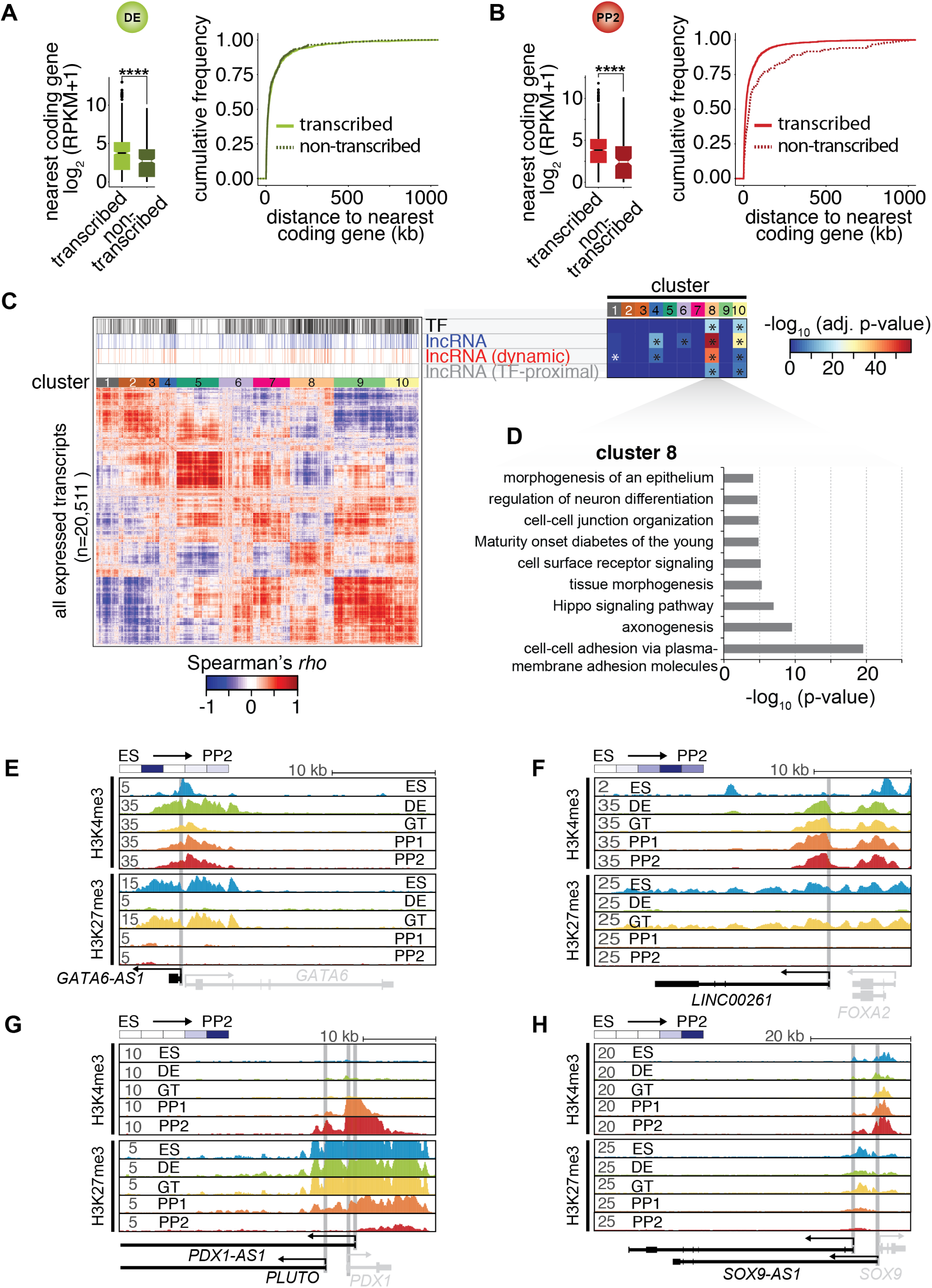
Related to Figure 1. Characterization of lncRNAs expressed during pancreatic differentiation. **(A**,**B)** Left: Expression of the single nearest coding genes (*±* 1000 kb) in *cis* to transcribed and non-transcribed lncRNAs at the DE stage (A) or PP2 stage (B). Log_2_ transformed mean expression values (RPKM + pseudocount) from two biological replicates were used to generate the box plots (****, p-value < 0.0001, Wilcoxon rank sum test). Right: Corresponding cumulative distance distribution functions. **(C)** Heatmap of the hierarchically clustered expression correlations (Spearman’s rho) of all RNAs transcribed during pancreatic differentiation (with RPKM ≥ 1 in at least ten out of 38 tissues). Transcription factor (TF)-encoding mRNAs, lncRNAs (all), dynamically expressed lncRNAs (RPKM ≤ 1 in at least one stage (ESC to PP2)), and TF-proximal lncRNAs are highlighted above the heatmap. Clusters 8 and 10 are significantly enriched for all of these RNAs (*, p-value < 0.03, Fisher test). **(D)** Gene ontology and KEGG pathway analysis for all coding genes in cluster 8 (p-value < 0.05, Fisher test). The full list of significantly enriched terms is shown in **Table S1C. (E-H)** H3K4me3 and H3K27me3 ChIP-seq tracks of loci containing lncRNAs *GATA6-AS1* (A), *LINC00261* (B), *PDX1-AS1*/*PLUTO* (C), or *SOX9-AS1* (D) during pancreatic differentiation of CyT49 hESCs.

**Figure S2.**
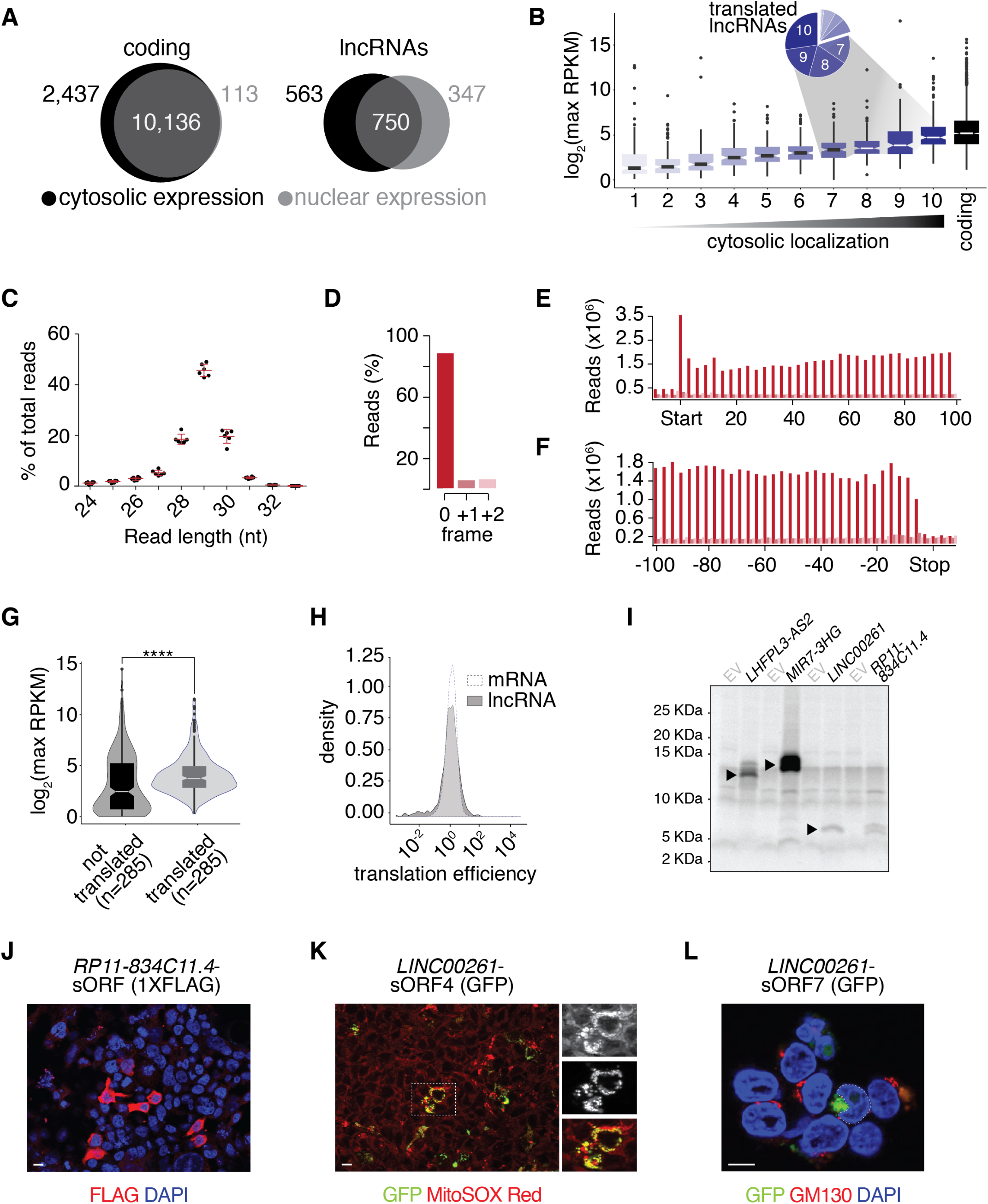
Related to Figure 2. Cytosolic lncRNAs engage with ribosomes. **(A)** Venn diagrams showing the number of coding RNAs (left) and lncRNAs (right) with RPKM ≥ 1 across two biological replicates in cytosolic and nuclear factions. **(B)** Box plots of maximum lncRNA expression (RPKM + pseudocount) across 38 tissues binned by their degree of cytosolic localization (measured as nuclear/cytosolic lncRNA expression ratio deciles in PP2); the expression of all PP2-transcribed coding RNAs with dynamic expression during differentiation is included for reference. The pie chart summarizes the proportions of translated lncRNAs within each cytoplasmic localization decile. **(C)** Read length distribution (nt) of Ribo-seq fragments across replicate Ribo-seq experiments (n = 6 biological replicates). **(D)** Position of the inferred P-sites of the ribosome footprints relative to the reading frame of PP2-transcribed coding genes. **(E-F)** Coverage of 29 nt footprint P-sites around the start codons (E) or stop codons (F) of PP2-transcribed coding genes. **(G)** Box plots comparing maximum expression of translated and untranslated lncRNAs (RPKM + pseudocount) across 38 tissues (****, p-value = 2.122×10-8, Wilcoxon rank sum test). For the untranslated set, 285 untranslated PP2-expressed lncRNAs were selected randomly. **(H)** Density plots comparing the translation efficiencies of PP2-expressed mRNAs and lncRNAs. **(I)** Autoradiograph of radiolabeled in vitro translation products derived from full-length *LHFPL3-AS2, MIR7-3HG, LINC00261*, and *RP11-834C11*.*4*. EV, empty vector. **(J)** Anti-FLAG immunofluorescence staining of HEK293T cells transiently transfected with a PGK-*RP11-834C11*.*4*-sORF-1xFLAG construct. **(K)** Microphotograph of HEK293T cells transiently transfected with a PGK-*LINC00261*-sORF4-GFP construct with mitochondria labeled by MitoSOX Red. **(L)** Golgi immunofluorescence staining (anti-GM130) of HEK293T cells transiently transfected with a PGK-*LINC00261*-sORF7-GFP construct. Scale bars = 10 µm.

**Figure S3.**
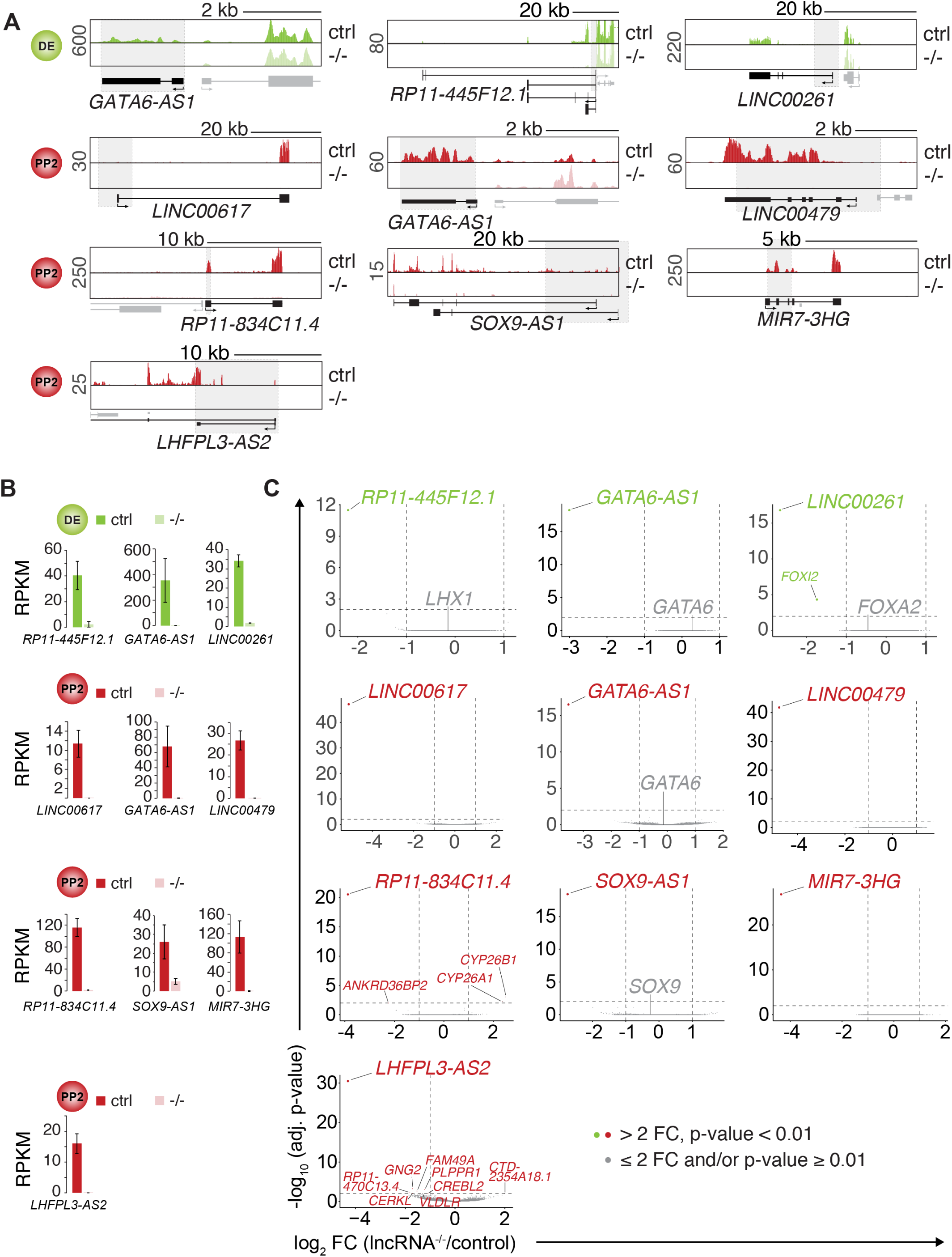
Related to Figure 3. Minor gene expression changes in definitive endoderm or pancreatic progenitor cells after lncRNA deletion. **(A)** Genome Browser snap shots of RNA-seq signal at the indicated lncRNA loci in control (ctrl) and lncRNA knockout (KO; -/-) DE (green tracks) and PP2 (red tracks) stage cells. Genomic deletions are indicated by gray boxes. **(B)** Bar graphs showing expression of indicated lncRNAs in control and lncRNA KO DE (green) and PP2 (red) cells quantified by RNA-seq. Data are shown as mean RPKM *±* S.D. (n = 2 independent differentiations of two independent KO clones, except for *SOX9-AS1* for which one clone was differentiated twice). **(C)** Volcano plots displaying gene expression changes in control versus lncRNA KO DE (green) or PP2 (red) cells. Differentially expressed genes (DESeq2; > 2-fold change (FC), adjusted p-value < 0.01; vertical and horizontal dashed lines indicate the thresholds; n = 2 independent differentiations of two independent KO clones, except for *SOX9-AS1* for which one clone was differentiated twice) are shown in green (DE) and red (PP2). TF genes in cis to deleted lncRNAs are shown in gray (gray dots represent genes with ≤ 2-fold change and/or adjusted p-value ≥ 0.01).

**Figure S4.**
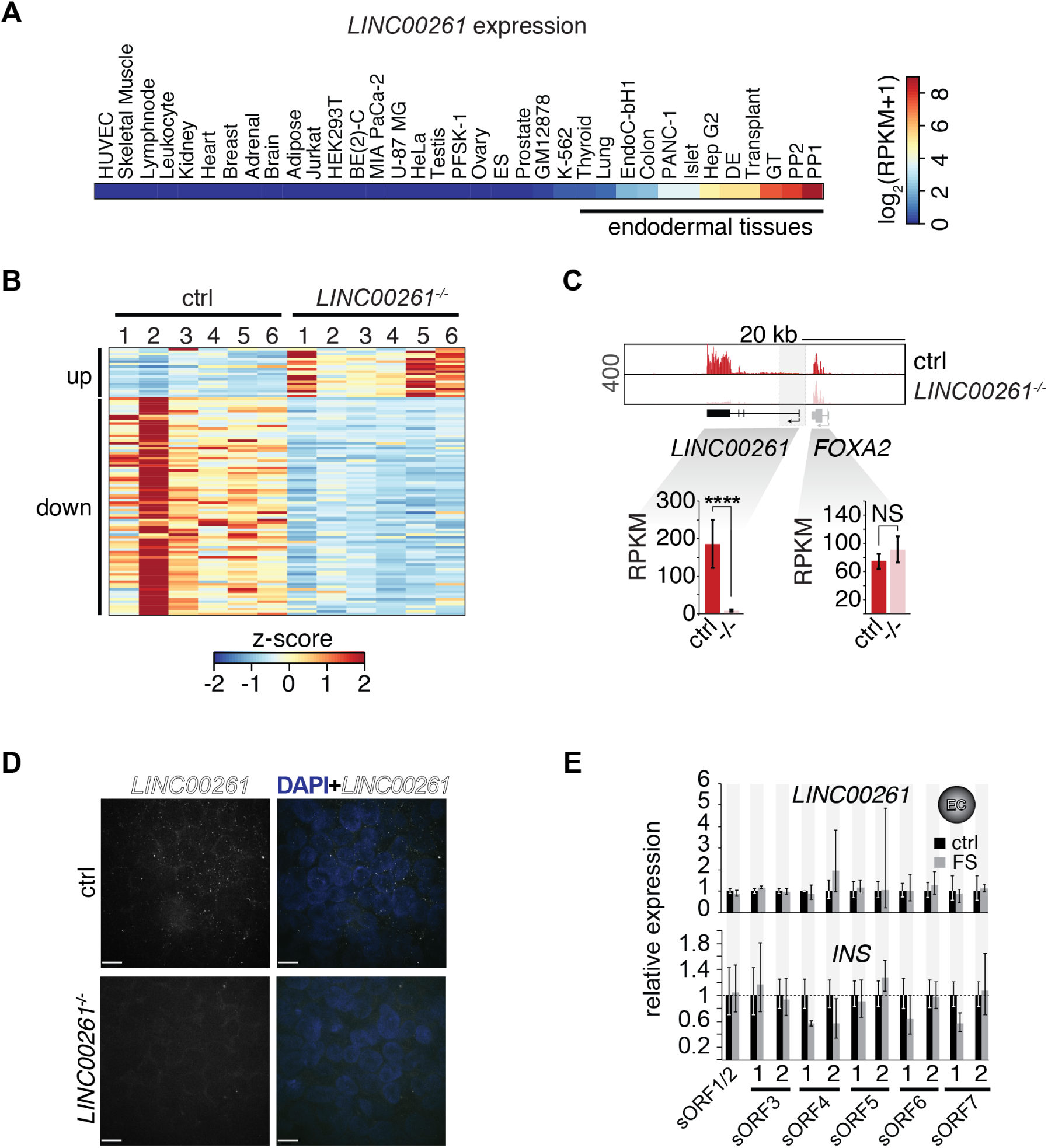
Related to Figure 4. Characterization of LINC00261-deleted pancreatic progenitor cells. **(A)** RNA-seq expression heatmap of *LINC00261* across 35 cell types/tissues originating from all three germ layers (shown as RPKM + pseudocount). **(B)** Heatmap showing K-means clustering of 108 differentially expressed genes (DESeq2; > 2-fold change (FC), adjusted p-value < 0.01) between PP2 cells from control (ctrl) and *LINC00261*^-/-^ H1 hESCs (based on expression z-score; n = 6 independent differentiations). **(C)** Top: Genome Browser snap shot of RNA-seq signal at the *LINC00261/FOXA2* locus in control and *LINC00261*^-/-^ PP2 stage cells. Genomic deletions are indicated by gray boxes. Bottom: Bar graphs showing *LINC00261* and *FOXA2* expression in control and *LINC00261*^-/-^ PP2 cells quantified by RNA-seq. Data are shown as mean RPKM *±* S.D. (n = 6 independent differentiations of four independent KO clones). ****, p-value < 0.0001; NS, p-value > 0.05; t-test. **(D)** LINC00261 smRNA FISH in control and *LINC00261*^-/-^ PP2 cells. Scale bars = 8 µm. **(E)** qRT-PCR analysis of *LINC00261* (top) and INS (bottom) expression in control and *LINC00261*-sORF-FS H1 hESC clones at the endocrine cell (EC) stage. Data are shown as mean *±* S.E.M. (n ≥ 3 independent differentiations for each clone).

